# Delineating Transdiagnostic Subtypes in Neurodevelopmental Disorders via Contrastive Graph Machine Learning of Brain Connectivity Patterns

**DOI:** 10.1101/2024.02.29.582790

**Authors:** Xuesong Wang, Kanhao Zhao, Lina Yao, Gregory A. Fonzo, Theodore D. Satterthwaite, Islem Rekik, Yu Zhang

## Abstract

Neurodevelopmental disorders, such as Attention Deficit/Hyperactivity Disorder (ADHD) and Autism Spectrum Disorder (ASD), are characterized by comorbidity and heterogeneity. Identifying distinct subtypes within these disorders can illuminate the underlying neurobiological and clinical characteristics, paving the way for more tailored treatments. We adopted a novel transdiagnostic approach across ADHD and ASD, using cutting-edge contrastive graph machine learning to determine subtypes based on brain network connectivity as revealed by resting-state functional magnetic resonance imaging. Our approach identified two generalizable subtypes characterized by robust and distinct functional connectivity patterns, prominently within the frontoparietal control network and the somatomotor network. These subtypes exhibited pronounced differences in major cognitive and behavioural measures. We further demonstrated the generalizability of these subtypes using data collected from independent study sites. Our data-driven approach provides a novel solution for parsing biological heterogeneity in neurodevelopmental disorders.

## Introduction

Attention deficit/hyperactivity disorder (ADHD) and autism spectrum disorder (ASD) are neurodevelopmental disorders that typically manifest in childhood, impact social function, and impair academic achievement^1,2^. ADHD across clinical subtypes is characterized by inattention, hyperactivity and impulsivity, while ASD presents with deficits in social communication and interaction accompanied by restricted, repetitive behaviors, and sensory symptoms^3^. Despite these seemingly independent core diagnostic criteria, there is a significant clinical overlap between ADHD and ASD^2,4^. Evidence from research indicates shared genetic risk factors and neuropsychological vulnerabilities, particularly in executive function and emotional recognition domains^2,5,6^. Adding to the complexity, both ADHD and ASD diagnoses, according to the Diagnostic and Statistical Manual of Mental Disorders (DSM) or the International Classification of Diseases, encompass a wide spectrum of symptom combinations. This symptom-based approach results in significant clinical heterogeneity, which aggregates patients with diverse symptoms, severity and mechanisms within a single medical diagnosis^7,8^. Beyond such clinical heterogeneity within each disorder, there is a significant co-morbidity of the two conditions: an estimated 30 to 50 % of individuals diagnosed with ASD also meet the criteria for ADHD^9^, and roughly 14% of the children with ADHD also have ASD^10^. By definition, co-morbidity suggests the co-occurrence of many diseases in any one patient may share similar causal mechanisms, which can not only exhibit distinct neurofunctional deficits but also complicate the treatments of patients^11^. Together, this combination of heterogeneity and co-morbidity complicates the task of discerning the underlying mechanisms of these disorders and devising effective interventions^12^. As such, there is an urgent need to identify subtypes within these broad clinical categories.

Research using functional magnetic resonance imaging (fMRI) to study brain functional connectivity may provide a promising avenue for identifying subtypes that explain biological heterogeneity in ADHD and ASD. Previous investigations have highlighted that resting-state fMRI connectivity involving the ventral attention network can predict treatment response in ADHD^13^ and assist in ASD diagnosis^14^. A meta-analysis further uncovered abnormalities in functional connections involving the frontal and posterior cingulate cortices from studies of group-level comparison between both ADHD and ASD with a healthy population^15^. Nonetheless, many studies have traditionally examined functional connectivity features independently (via vectorization), overlooking the potential benefits of considering the connectivity topology in understanding disorder vulnerability and resilience^16^. In response to this gap, some research efforts have shifted to analyze high-level brain graph features, such as participation coefficient and connectedness coefficient derived from the binarized functional connectivity graph^17,18^. Yet, focusing solely on these graph features may overlook essential global brain network properties. To utilize the complete brain graph information for downstream brain dysfunction analysis, deep learning-based end-to-end graph convolutional networks (GCNs) offer substantial advances in brain network representation, holistic mapping, diagnosis and generation^19^. Increasing numbers of research studies have shown the promising diagnostic ability of GCNs for ADHD and ASD^20,21^. Nevertheless, most of the existing studies have been conducted under a case-control framework by examining the differences between a single disease and healthy controls, overlooking the considerable heterogeneity and comorbidity inherent in ADHD and ASD^22^.

Building on the hypothesis that ADHD and ASD present varying manifestations of an overarching neurodevelopmental disorder^23^, our objective was to uncover novel transdiagnostic subtypes within these diagnoses. To this end, we employed graph neural networks to extract diagnostic-independent representations shared across both disorders. Recent advancements in self-supervised learning, especially contrastive learning, have paved the way to discern patterns from high dimensional functional connectivity data without being anchored to clinical diagnoses^24,25^. Unlike supervised learning models that optimize parameters based on categorical clinical labels, contrastive learning establishes a metric that enhances similarities between two functional connectivity representations derived from the same individual while fostering contrasts between representations from various individuals. Integrating contrastive learning with GCNs has the potential to delineate shared functional connectivity patterns between ADHD and ASD, while also parsing heterogeneity within each clinical diagnostic group. Accordingly, we developed a contrastive graph convolutional model to extract the ADHD and ASD representative graph features from brain functional connectivity networks. Our modelling framework learns disease-agnostic representations, which then serves as the foundation for building a clinical population graph spanning both ADHD and ASD. Using spectral clustering on the graph data, we were able to delineate distinct transdiagnostic subtypes. With three large-scale resting-state fMRI cohorts, including ADHD-200^26^, Autism Brain Imaging Data Exchage (ABIDE)-I and ABIDE-II^27,28^, our approach successfully identified two robust subtypes spanning both ADHD and ASD. Furthermore, we evaluated differences in independent functional and clinical measures that were not used in subtype identification. We also ensured the generalizability of our findings by conducting subtyping analyses on a separate, held-out dataset. Collectively, our approach seeks to fully harness the brain graph information, revealing novel and insightful transdiagnostic subtypes to disentangle the heterogeneity and comorbidity of ADHD and ASD. In doing so, we provided initial evidence for the neuropathological categorization of neurodevelopmental disorders, thus advancing the development of precision mental health.

## Results

We first conducted the contrastive brain graph analysis and subtype identification using composite discovery data and then validated our results using held-out data. As we resolve to analyze the transdiagnostic relationships, we strategically combined two multi-sites datasets that cover ASD or ADHD patients for discovery analysis. The composite discovery data (including 268 ADHD, 531 ASD, and 881 neurotypical controls) combines a subset of ADHD-200 and the whole ABIDE-I. This subset of ADHD patients was selected from three sites with relatively balanced patient-to-healthy-control ratios. The ratio of training-validation-testing set was 7:1:2 and we tested the average model performance by randomly splitting dataset using six random seeds. The subtyping algorithm was trained on the training subset of the discovery set (N=1176), and the best model was saved by evaluation on the validation subset (N=168) and tested on the testing subset (N=336), while the subtype labels were assigned to all of the train-validation-test subsets. Such process was implemented on the discovery set (ABIDE-I and ADHD-discovery set). To further verify the transferability of the acquired subtypes, we introduced a held-out dataset (including 45 ADHD, 453 ASD, and 744 neurotypical controls) which contains the remaining site patients of ADHD-200 and the whole ABIDE-II dataset. This composite held-out dataset was employed for prediction rather than training the subtyping algorithm (see Participants section for more details).

### Contrastive functional connectivity representation defines two subtypes across ADHD and ASD

In order to learn generalizable subtypes across ASD and ADHD, the proposed model should focus on transdiagnostic features rather than clinical labels. In light of this, our proposed contrastive graph learning approach (see Supplementary Figure 1 and section Methods) was applied to brain graphs constructed using functional connectivity together with personal characteristic data (PCD) on the clinical population. The functional connectivity was extracted using Schaefer parcellation of 100 regions of interest (ROIs). Site effects were mitigated by applying the well-established statistical harmonization tool ComBat ^29^ to functional connectivity inputs meanwhile preserving biological variability such as age, sex, handedness, and three IQ measures. The extraction of graph representations was based on self-resemblance of two non-overlapping fMRI series acquired for the same subject, meanwhile maximizing intra-subject contrast regardless of their clinical labels. Neurotypical controls were also utilized to guide the learning of disease-specific contrastive graph representation. We then employed the resulting representations to induce a population graph whose adjacency matrix was computed using similarities between subject features. Spectral clustering along the population graph was further conducted to identify transdiagnostic subtypes. As the graph representation was learnt in a diagnosis-agnostic manner, the resulting subtypes can be applicable to both ASD and ADHD subjects.

Our approach discovered two distinct subtypes across ADHD and ASD populations(Figure 1). where the optimal number of the subtypes was suggested by the highest Calinski-Harabasz index for the clustering algorithm (Supplementary Figure 2). To understand which neural circuits are crucial for identifying the two subtypes, we examined connectivity differences in terms of gradient-based feature weights (see section Methods for more details of the feature weights calculation). Gradient-based methods were originally proposed to improve interpretability of deep learning methods^30^. For instance, recent work^31^ computed the partial derivatives of the Austism labels with respect to the individual functional connectivity links from the brain connectom to identify discriminative functional connections. Many studies have emerged since to adopt gradient-based methods for brain imaging analysis^32–34^. We hereby compute the gradients of our subtyping model predictions to individual connectivity and group the resulting connectivity feature weights by subtyping labels. The resulting mean connectivity feature weights (Figure 2) revealed critical neural circuits that contributed to defining each of the two subtypes. As the number of the identified subtypes is 2, the gradient weight for one connectivity would be positive for one subtype and negative for the other subtype, indicating such connectivity is contributing positively to one subtype and has a negative contribution to identifying the other subtype. Hence, the resulting feature weight map on two subtypes are inversely correlated. The neural circuit pattern of subtype 1 mainly involved the somatomotor network (SMN), visual network (VN), ventral attention network (VAN), and dorsal attention network (DAN). Both the inter- and intra-network connections of these networks have acquired large feature weights (Figure 2 a, b, c). By contrast, the neural circuit pattern of subtype 2 was dominated by both inter- and intra-network connections involving the default mode network and frontoparietal control network (Figure 2 a, b, c). As the model is capable of predicting the individual connectivity feature weights, we further examined the subtype differences by computing two-sample t-tests on the where the feature weights of one subject was treated as a sample in a subtype group. Such comparison can provide a reliable measure as opposed to the simple difference of the averaged feature weights of two subtypes. Our results showed that connectivity gradients captured significant differences in the crucial networks of subtype 1 including SMN, VN, VAN, and DAN (Figure 4 a), which were not identifiable using raw connectivity features (Figure 4 b). The mean FC inputs show great resemblance (Supplementary Figure 4), which justifies the infeasibility of detecting subtypes on raw connectivity features.

**Figure 1.**
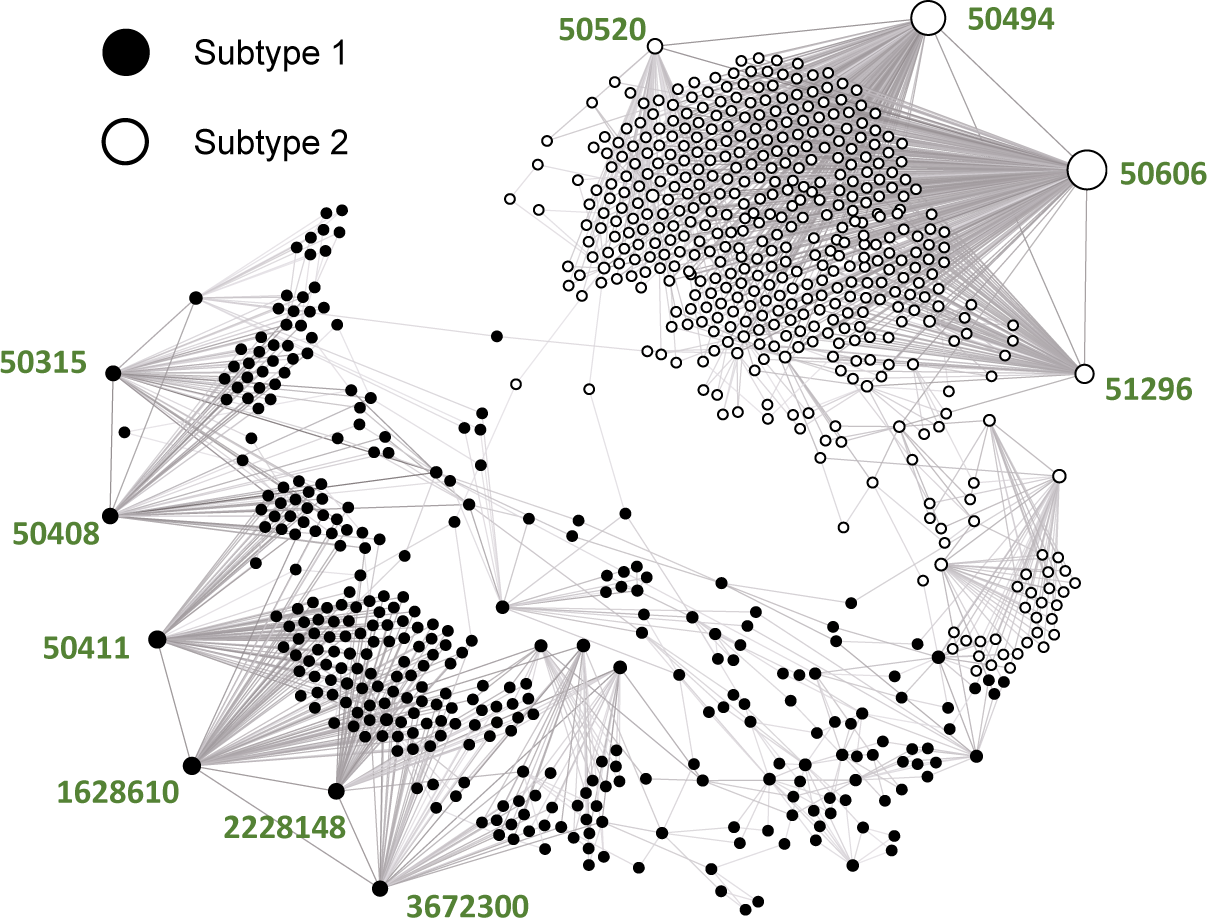
Contrastive functional connectivity representation identifies two distinct subtypes across ADHD and ASD. Each node represents an individual in the ADHD and ASD population, whereas the edge connecting two individuals indicates strong feature similarities between their contrastive features extracted by our proposed method. The node size depicts the degree of each node and the node colors differentiate the subtype labels. The colored number indexes represent the subject names of those “hub patients” who share the most feature similarities with other individuals.

**Figure 2.**
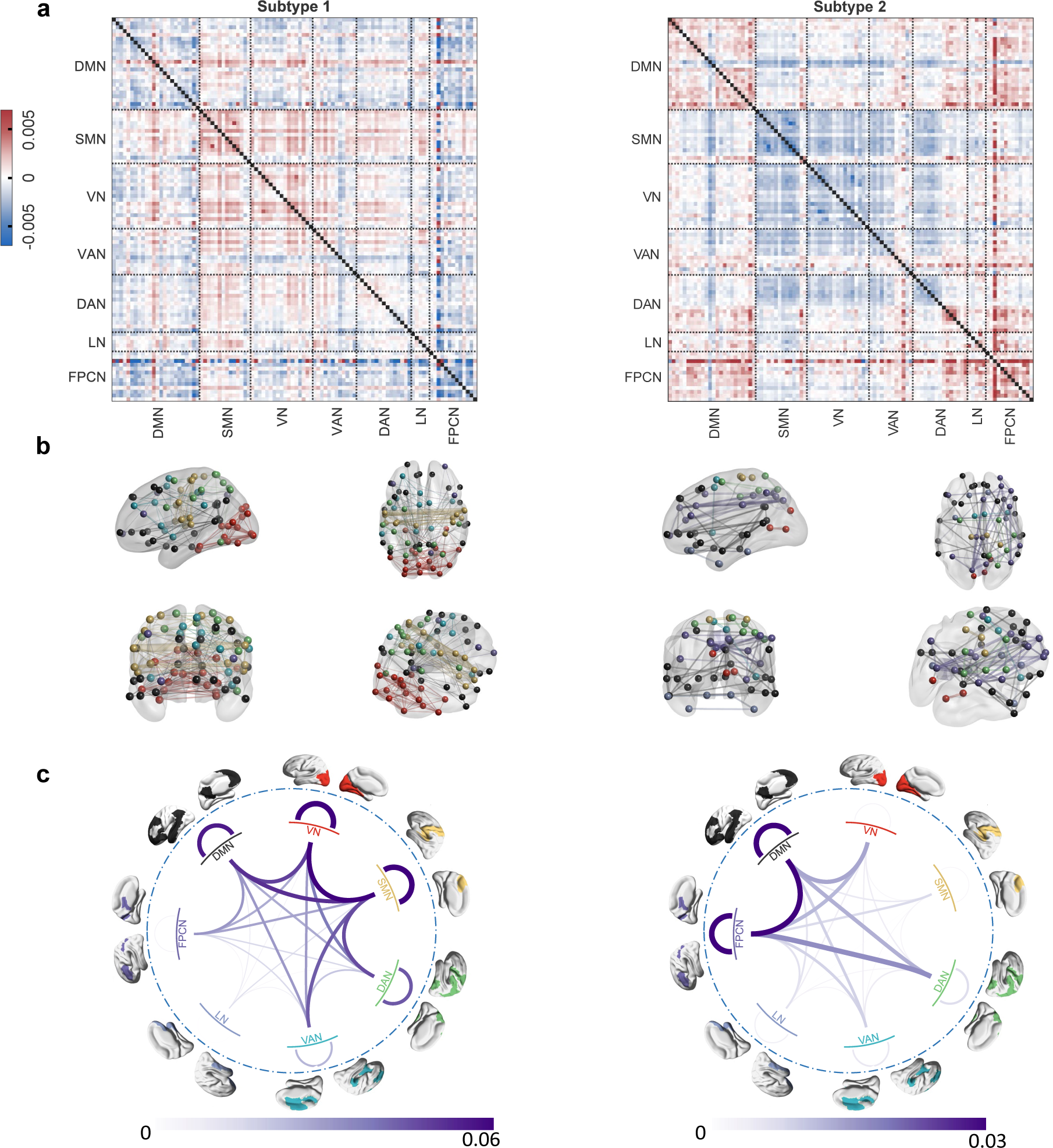
Mean gradient-based connectivity feature weights of the two subtypes on the transdiagnostic discovery set comprised of ABIDE-I and the ADHD-discovery set. The gradients are obtained from a subtyping classifier trained on the observed subtypes as ground truth, **a.** Mean connectivity gradient matrices where a positive gradient indicates the positive contribution of a connectivity feature to the subtype. The ROIs are reordered by seven brain networks: DMN: default mode network; SMN: somatomotor network; VN: visual network; VAN: ventral attention network; DAN: dorsal attention network; LN: limbic network; FPCN: frontoparietal control network. **b.** Visualization of connectivity patterns on the brain. Each ROI is represented by a node whose color was decided by the corresponding network. The edge width indicates absolute gradient value for a specific connectivity. The edges with strength below the 0.03 *×* maximum value are hidden for visual clarity. **c.** Inter- and Intra connectivity gradients on seven brain networks. The cord width describes the network-level connectivity gradients. The network color matched with the visualization on the brain.

To explore whether our subtyping method can be used to identify similar subtypes in individual diagnoses, we evaluated our analysis pipeline on ABIDE-I and the ADHD-discovery set separately. The experiment aims to test whether both subtypes can be identified separately in ASD and ADHD. Again, two subtypes were discovered from each dataset (Supplementary Figure 5 a,b). We quantified the similarity between these subtypes by computing Pearson correlation coefficients between the individual connectivity feature weights of each subtype and the weights of two transdiagnostic subtypes (see Supplementary Table 2). One of subtypes obtained from the individual ABIDE-I dataset suggested moderate resemblance towards transdiagnostic subtype 1 with a significant correlation coefficient of 0.34 (Supplementary Table 2 ABIDE-I subtype 2) and emphasis on connectivities within SMN, VN, and DAN (Supplementary Figure 5 a). For the ADHD-discovery set, both identified subtypes from this individual set (Supplementary Figure 5 b) showcased strong subtype similarities with the corresponding transdiagnostic subtypes with high correlation coefficient (*r* = 0.58, *p* < 10*^−^*^5^ for subtype 1 and *r* = 0.66, *p* < 10*^−^*^5^ for subtype 2 (see Supplementary Table 2 ADHD discovery set subtype 1 and 2). The crucial networks discovered by ADHD-discovery set subtype 1 are SMN, VN, DAN, while the subtype 2 showed distinct connectivity patterns in DMN and FPCN. Both echo the conclusions obtained on the transdiagnostic subtypes.

To further validate whether our model can be used for identifying resemblant subtypes from unseen patients with ASD or ADHD, we further tested our pipelines on held-out data comprising ABIDE-II and ADHD-validation set. The resulting FC gradients of both subtypes in ABIDE-II (Figure 3) are highly correlated with the corresponding subtypes in the transdiagostic discovery set with positive Pearson coefficients (*r* = 0.54, *p* < 10*^−^*^5^ for subtype 1 and *r* = 0.48, *p* < 10*^−^*^5^ for subtype 2, see Supplementary Table 2 ABIDE-II results), whereas the gradient of the ADHD-validation set did not support such conclusion (Supplementary Table 2 ADHD-validation set results). Such divergence may be attributed to the small sample size since only 45 patients are available from the ADHD-validation set. Overall, transdiagnostic subtype 2 shared higher correlation coefficients on the held-out set, which showed distinct patterns on DMN and FPCN.

**Figure 3.**
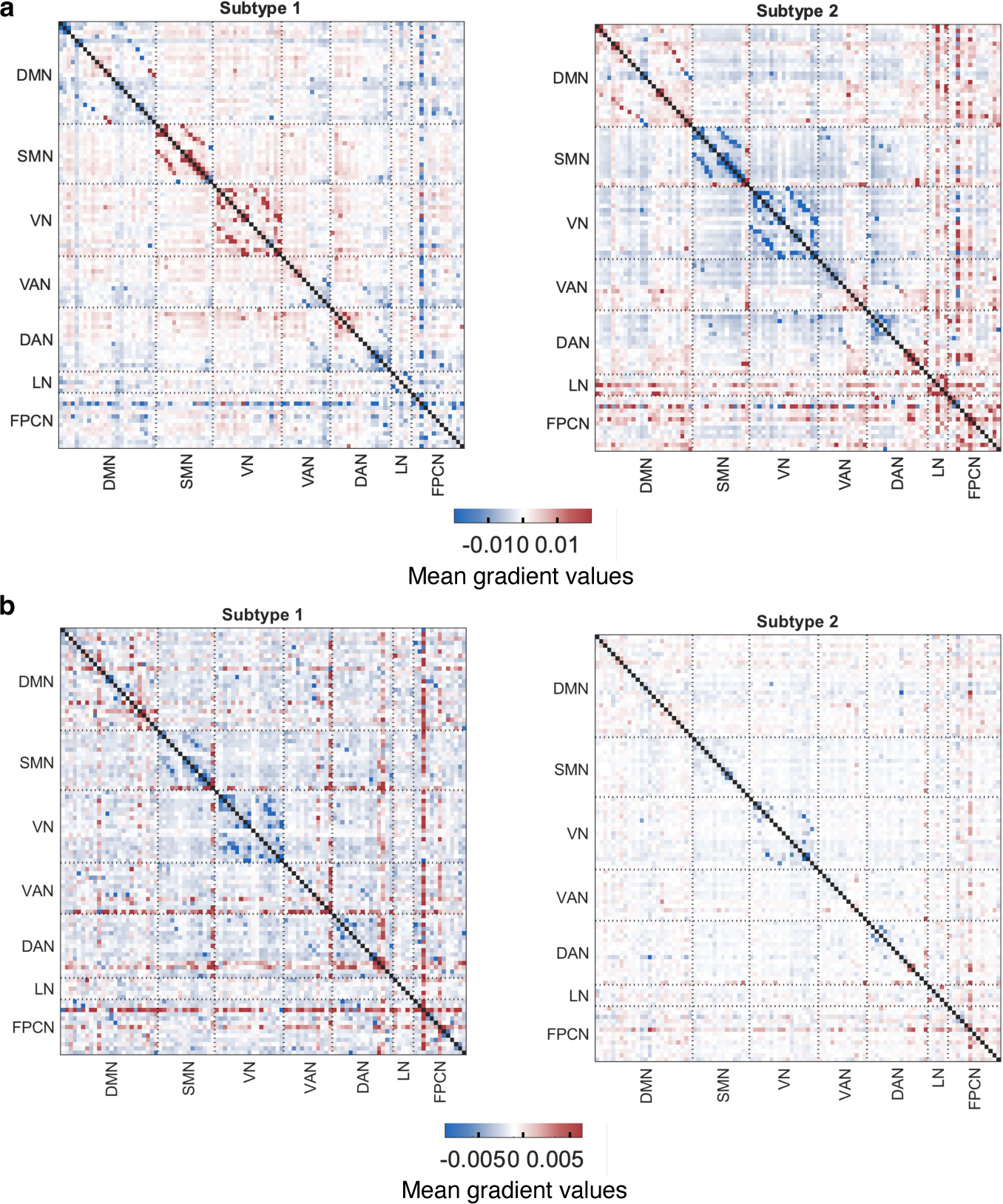
Connectivity gradients of two subtypes in individual replication sets. The gradients are obtained by retyping on two replication sets separately. Positive gradients indicate a positive contribution of connections to the subtype, while negative gradients signify a negative contribution **a**. Mean connectivity gradients of the two subtypes on ABIDE-II. **b.** Mean connectivity gradients of the two subtypes on the ADHD-validation set.

**Figure 4.**
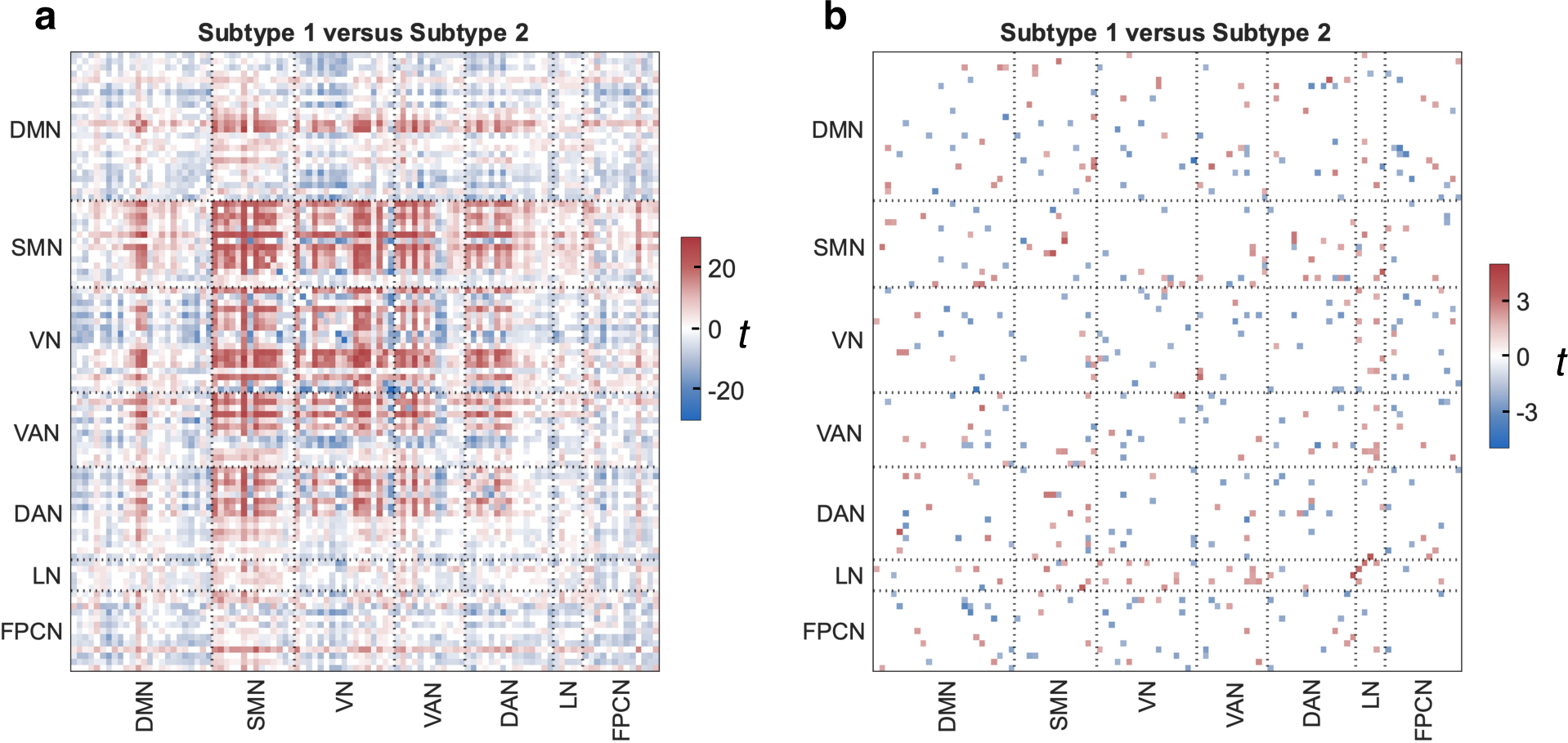
Connectivity differences between the two subtypes detected by independent t-test (FDR-corrected P < 0.05.a. Significant difference of gradient of raw functional connections (partial correlation). **b.** Significant difference of raw connectivity features.

### Subtype differences on demographic, cognitive and behavioral data

We further examined differences between identified subtypes in demographic, cognitive and behavioral measures. Three demographic features between the two diseases were firstly compared: age, gender, handedness (Supplementary Figure 6 a,b). Subtype 1 is younger than subtype 2 with significant difference(Supplementary Table 1). We then investigated differences between subtypes in the cognitive and behavioral assessments for the discovery set. Three transdiagnostic behaviour measures are included as training features: full-scale intelligence quotient (FIQ), verbal intelligence quotient (VIQ), and performance intelligence quotient (PIQ). Patients in subtype 2 achieved higher scores on these IQs (see Supplementary Table 1, 5, 4). Several disease-specific behavioral measures were also compared between subtypes. For the ADHD-discovery set, the two subtypes behave similarly with *p* > 0.05 on behavioral scores including ADHD index, inattentive and hyperactivity / impulsivity scores and their medical status (Supplementary Table 5). For the ABIDE-I dataset, subtype 1 constitutes the major ASD and Pervasive Developmental Disorder-Not Otherwise Specified (PDD-NOS) patients while in subtype 2 the dominant patients are Aspergers and Aspergers or PDD-NOS groups (Supplementary Table 4). Within ABIDE-I, subtype 2 showed significantly lower scores on three ASD clinical measures: Abnormality of Development Evident at or Before 36 Months (D) Total (ADI_R_ONSET_TOTAL_D), Classic Total Autism Diagnostic Observation Schedule Score (Communication subscore + Social Interaction subscore, ADOS_TOTAL) and Social Total Subscore of the Classic ADOS (ADOS_SOCIAL) (Supplementary Table 4). Subtype 2 also achieved significant higher scores on most of cognitive measures (starting with WISC_*, Wechsler Intelligence Scale for Children) except for WISC_IV coding scaled.

We further examined the demographic, cognitive and behavioral results on the held-out datasets (ABIDE-II and the ADHD-validation set). For ABIDE-II, subtype 1 has significant younger populations which echoed the conclusion of the transdiagnostic discovery set (Supplementary Table 6). The results from the DSM_IV measurements were also consistent with the discovery set where significant differences are observed between subtype 1 and 2 with more ASD participants in subtype 1, and higher Aspergers proportion in subtype 2. Similar to the discovery set, significant differences were recognized on FIQ and VIQ but not PIQ (Supplementary Table 6). Three clinical measurements on the held-out datasets also echoed the conclusion of the discovery set (ABIDE-I) where subtype 1 achieved higher scores: ADI_R_ONSET_TOTAL_D, ADOS_G_TOTAL, and ADOS_G_SOCIAL. Surprisingly, we discovered that subtype 1 has achieved significant higher scores on two ADHD-related clinical measurements: CBCL 6-18 Attention Problems T Score and CBCL 6-18 Attention Deficit/Hyperactivity Problems T Score.

In order to verify whether subtypes defined by the connectivity features can be identified using the demographic, cognitive and behaviour features alone, we ran a K-means clustering algorithm using the following six features that are transdiagnostic between ADHD and ASD: gender, handedness, age and three IQs: FIQ, VIQ, and PIQ (See Supplementary Figure 7 a, b). While these features shared an high accuracy on the ADHD-discovery set, the validation results on the ADHD-replication set and ABIDE-II are only 60% and 66% respectively as apposed to our results of 68.9% and 79.9%, suggesting that the subtypes acquired by PCD features were less transferable and replicable than our contrastive features.

### ASD and ADHD population graph investigation

As the node sizes depict the node degree in the population graph (Figure 1) and the edges represent strong subject similarities, a large node with many connecting edges suggested that this particular patient showed resemblance with many other patients on contrastive features and therefore their subtype labels would affect the results for most of the patients in the population graph. We therefore investigated whether the “hub patients” functional connectivity, demographic or biological features contribute to the subtyping results. We sorted the patients by their node degree and selected the top-10 hubs in both subtypes. Most Subtype 2 patients were linked to four hubs from ABIDE-I: 50606, 50494, 50520, 51296 (subject id), whereas Subtype 1 patients are connected by more scattered hubs from both ADHD-discovery set and ABIDE-I patients: 1628610, 3672300, 2228148 from ADHD-discovery set and 50411, 50408, 50315 from ABIDE-I. We first placed the demographic and behavioral features of these hubs into the corresponding distributions of the non-hub patients and evaluated their likelihood (Supplementary Figure 8). Two patients 50606 and 50494 in subtype 2 were identified as outliers, since either their ages were substantially older or they achieved much lower IQ scores. We also compared hub patients’ raw FC inputs with the average connectivities of non-hub patients in the corresponding subtype (Supplementary Figure 9). Most of the hub patients were found to have marginally higher connectivities on the visual networks or somato-motor networks than non-hub patients.

### Validation of the subtype transferability across datasets

To investigate whether the subtypes acquired from the transdiagnostic population are transferable to unseen patients, we assigned labels for new individuals with two solutions: either by choosing the subtype of the most similar subject in the transdiagnostic population as the prediction or prediction using a classifier trained by regarding transdiagnostic subtypes as ground truth. First, we applied the same subtyping pipeline to individual diagnoses and map their subtyping results to the transdiagnostic discovery set (see Supplementary Figure 3). For subtypes acquired from ADHD alone, patients in ASD in the transdiagostic popultion were unseen by the model and therefore can be used for validation purpose. Both subtypes from the transdiagnostic discovery set were recognized in the individual diseases. To assess the robustness of the obtained subtypes, we conducted several transferability experiments on individual discovery set (ABIDE-I and ADHD-discovery set) as well as replication set (ABIDE-II + ADHD-validation set) (see Figure 5). We initially evaluated the transferability from the transdiagnostic discovery set to individual discovery sets. The subtypes obtained from the trans-discovery set (Figure 1) were utilized as predictions and the ground truth subtypes were acquired from training and testing on individual discovery sets, i.e., ABIDE-I, or ADHD-discovery set. The result on the ADHD-discovery set achieved a higher accuracy of 85.8% compared with 79.7 % on the ABIDE-I, which indicates that each subtype can be identified in individual diagnoses with high transferable accuracy (see the confusion matrix in Figure 5 a).

**Figure 5.**
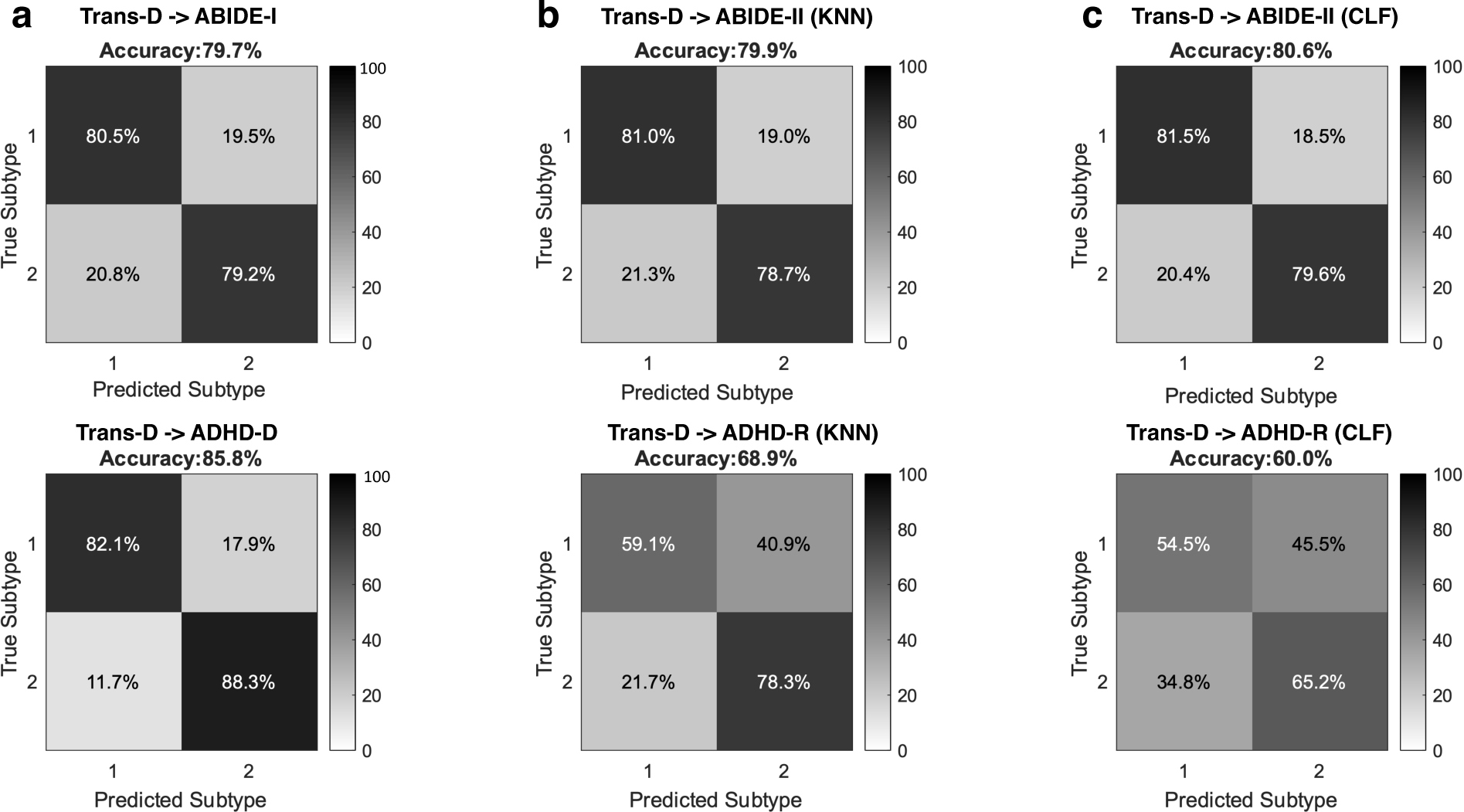
Confusion matrix for model transferability. Subtypes obtained from the transdiagnostic discovery set (Trans-D) are utilized to predict the subtypes on the individual discovery set and replication set. **a.** Confusion matrix from Trans-D to ABIDE-I and ADHD-discovery set (ADHD-D). We run the same pipeline on the individual disease and obtained their subtypes as ground truths and the subtypes from the transdiagnostic population are treated as predictions. **b.** Confusion matrix on the replication dataset of ABIDE-II and ADHD-validation set (ADHD-R) using KNN-based method. We run the same pipeline on the replication set to obtain subject embeddings, the subtyping predictions are then assigned based on their neighbours in the trans-D set. The ground truth is obtained by running the pipeline on the replication set separately. **c.** Confusion matrix on ABIDE-II and ADHD-validation set using a subtyping CLassiFier (CLF) which was trained using the subtyping labels from the Trans-D set.

Additionally, we validated the subtype findings using ABIDE-II and ADHD-validation set to confirm the replicability of the identified subtypes. The analysis was implemented from two perspectives, KNN-based method and subtyping classifier. KNN-based method employed the trans-discovery subjects as neighbours to label subtypes of the unseen patients, whereas subtyping classifier explicitly train a classifier on the trans-discovery labels to predict subtyping labels for new patients. Transferability is more robust in ABIDE-II than in ADHD-validation set, where the average accuracy for ABIDE-II was 79.9% for KNN method (Figure 5 b) and 80.6% for subtyping classifiers (Figure 5 c), and the average accuracy on ADHD-validation set was 68.9% for KNN-based method (Figure 5 b) and 60% for classifiers (Figure 5 c). Such difference may be attributable to the small dataset size of ADHD that consequently may generate less reliable subtyping labels.

## Discussion

Heterogeneity and comorbidity have been frequently reported in ADHD^35–37^ and ASD^38^, which precludes delivery of effective and customized treatment to distinct subgroups within the same clinical diagnosis. In light of this, we have developed a novel subtyping framework utilizing contrastive graph machine learning, which successfully identified two transdiagnostic subtypes characterized by differing functional connectivity gradients, and cognitive and behavioral profiles. In particular, subtype 1 exhibited significantly increased FC gradients in somatomotor network, visual network, ventral attention network and dorsal attention network. Subtype 1 also showed significantly lower cognitive scores (including FIQ, VIQ and PIQ) than subtype 2 and healthy controls. For ASD patients in subtype 1, they displayed higher total ADOS scores, where an ADOS score ranging from 30-36 suggests mild to moderate autism, and a score greater than 37 is classified as severe autism^39^. Subtype 1 patients also achieved lower cognitive behaviour scores with the WISC assessments including Verbal Comprehension Index (VCI), Perceptual Reasoning Index(PRI), Working Memory Index (WMI) and the Processing Speed Index (PSI), where a lower score below 90 suggests below average intellectual ability. In contrast, subtype 2 exhibited significantly increased FC gradients in the default mode network and frontoparietal control network. Subtype 2 displayed clinical profiles that resembled those of healthy controls. Furthermore, the validation of these subtypes on the independent dataset ABIDE-II and ADHD200 indicated high transferability across datasets. In particular, both subtypes showed consistent functional connectivity gradients and intellectual capacity measurements, such as FIQ and VIQ, between the discovery data and held-out data. These findings provide further support for underlying neurobiological mechanisms that may contribute to substantial comorbidity between ASD and ADHD. These insights have the potential to facilitate future precision treatment efforts through stratifying individuals by subgroups across diseases that have more homogeneous characteristics, which may, in turn, improve treatment outcomes if specific treatments are better suited to a particular subtype. This moves the field going beyond reliance solely on clinical diagnoses and trial-and-error treatment approaches.

Our findings provide valuable insight into the heterogeneity of functional connectivity observed in individuals with ASD and ADHD, confirming previous research and highlighting distinct neural circuit patterns associated with specific subtypes. For ASD, our identified subtype 2 confirms previous evidence of brain dysfunction in DMN^40–42^. A study^40^ associated DMN with the main social deficit in children with ASD and identified fundamental aspects of posteromedial cortical heterogeneity. Interestingly, two recent studies of functional connectivity heterogeneity in ASD have shown very similar subtypes to ours ^43, 44^. Other work ^43^ detected three subtypes in a combined dataset, including ABIDE-II. Similar to this study, their important connectivities in subtype 1 and subtype 2 are almost complementary with subtle differences. Regarding ADHD, we found supporting evidence from various studies that pinpoint highly related regions^45–52^. These studies have consistently identified functional connectivities within and between DMN and FPCN, which is consistent with the patterns observed in our subtype 2. One study ^45^ also suggests hypoconnectivity in VAN, DAN, and SMN, which we observed in our subtype 1. There is also an increasing trend to parse heterogeneity within the ADHD diagnosis. For instance, three discrete subtypes are identified by Damien A 2015^53^ on a group of children with and without ADHD. Atypicality in DMN is discovered in all three subtypes. Their subtype 1 is specifically connected to regions of DAN and VN which parallels our subtype 1. Altogether, these findings justify the importance of pursuing subtyping efforts on the ADHD and autism diagnoses and converge with the findings in this investigation regarding the distinct networks or regions associated with specific subtypes.

Emerging evidence points to a shared behavioral dysfunction between ADHD and ASD, as highlighted by various studies^54–56^. Our observations regarding FPCN dysfunction in subtype 2 find support from several significant research efforts. For instance, literature ^6^ posited that the impairment of executive function could be a shared root for neurodevelopmental disorders like ADHD and ASD. Additionally, a subsequent study^57^ highlighted that executive function characteristics not only explain the co-occurrence of ADHD and ASD, but also underscore the intrinsic heterogeneity across these disorders, thereby hypothesizing subgroups with distinct mechanisms across disorders. This research delineated differences between subtypes and neurotypical controls, particularly in VAN and SMN, which corroborates our findings in subtype 1. While distinctions based on hyperactive/inattention symptoms were discernible, no functional connectivity differences were identified between their subtypes, implying a more complex mechanism underlying these disorders. We surmise that this might stem from the emphasis on direct functional connectivity measurements, rather than on connectivity gradients - an approach that our results affirmed as essential for discerning key connectivity features. In a separate endeavor, Lake et. al.^58^ confirmed the significance of FPCN and also pinpointed an accentuated hubness in VN. Furthermore, they identified overlapping domains in representation/recognition, and language or semantics across these disorders. Notably, literature^54^ detected shared classification markers between ASD and ADHD across LN, DMN, VAN, and VN. In sum, these insights pave the way for a more comprehensive understanding of both the inherent comorbidity shared aspects between ADHD and ASD.

To assess the association between transdiagnostic subgroups, demographic data, and behavior scores, we examined shared features spanning age, gender, handedness, FIQ, VIQ, and PIQ across diseases. Additionally, disease-specific scores were evaluated to elucidate subtype distinctions in the brain. Intriguingly, neither subtype predominated in ADHD or ASD, suggesting our identification of two genuinely transdiagnostic subgroups. Our age-related findings converge with literature ^59^ reporting that younger ASD individuals exhibited greater impairment in lateral temporal (a component of VN) and frontal (a part of FPCN) regions. This aligns with the FC gradients we observed in subtype 1. These effects might stem from enhanced neurogenesis, reduced synaptic pruning, diminished neuronal cell death, or abnormal myelination, as noted in literature^60^. Furthermore, another study^59^ established a link between FIQ values and CTs, where individuals with lower FIQ (akin to subtype 1 in our study) exhibited abnormalities in frontal (VAN, DAN), temporal (VN and part of SMN), and occipital (VN, SMN) cortices.

Regarding the disease-specific symptoms, both subtypes presented similarly in terms of the ADHD index, inattention, and hyper-impulsivity scores. We did not observe the distinct patterns suggested in literature^57^ for inattention and hyperactivity/impulsivity scores. In contrast, these subtypes showed varied results for ASD clinical diagnoses and behaviour scores. Our analysis suggests the model might capture a broader variance of ASD symptoms compared to ADHD. Specifically, subtype 1 comprises a higher percentage of ASD patients, while subtype 2 predominantly includes Asperger’s patients. This observation aligns with Lotspeich (2004)^61^ which posited that Asperger’s syndrome manifests milder neuroanatomical deviations compared to lower IQ individuals (as seen with ASD patients in our subtype 1). Additionally, the Asperger’s subgroup showed fewer abnormalities in Autism-related scores. This was echoed by Anderson (2011)^42^, which identified significant correlations between individuals with ASD and behavioral measures such as ADOS_SOCIAL, ADOS_COMM, the Social Responsiveness Scale (SRS), and VIQ. These scores depict verbal language deficits and deficits in social interaction. Another study^62^ found an association between atypical FC patterns and clinical severity in ASD, pinpointing regions similar to our findings. They revealed that heightened clinical severity was linked to enhanced connectivity regions within DMN such as the angular gyrus. In contrast, connectivity diminished in areas vital for multimodal integration like the superior frontal and parietal zones of the DAN. Additionally, subtype 2, including primarily Asperger’s patients achieved higher scores in IQ and verbal assessments, which showcased the distinction of language skills between Autism and Asperger’s syndrome. We also observed a subtype distinction on the Child Behaviour Checklist attention problems subscale (CBCL_6-18) where subtype 2 presented fewer anomalies than subtype 1. A similar study^63^ conducted subtyping on a comorbid group of ADHD and ASD patients and found that healthy controls exhibited fewer attention problems relative to both diseases. No discernible difference was detected between the two diagnostic groups, lending further weight to the notion of shared patterns across these disorders.

Several limitations and potential avenues for future exploration emerge from the current study. By selecting multiple scans or segmenting the ROI time series into non-overlapping windows, we created multiple views of the subjects. Such an approach may limit model flexibility, especially when additional scans are unavailable, or the data length is insufficient for segmentation.

Future studies might consider graph topology augmentation as a strategy to generate positive pairs for contrastive learning^64^. Furthermore, integrating behavioral and cognitive measures as training features could lead to biased feature importance on them and consequently weakened the effects of functional connectivity. Solely focusing on functional connectivity in subtyping analyses might overcome such limitations. Finally, given the limitations of data acquired in the same samples utilized here to represent neurodevelopmental disorders, we were unable to assess the treatment effects on the various subgroups or detail their responses, which worth future studies with data from clinical trials.

In summary, our study developed a contrastive graph representation learning framework to identify transdiagnostic subtypes across ADHD and ASD based on rs-fMRI functional connectivity. Our comprehensive analyses revealed two replicable subtypes that showed distinct functional connectivity gradients and associations with intrinsic large-scale brain networks. The neurobiological heterogeneity and comorbidity across the two diseases were underscored by subtype validation on held-out data. These subtypes further demonstrated differences in demographic characteristics and clinical severity. Additionally, both subgroups exhibited diverging patterns of intellectual performance for the ASD patients. Our subtype discovery efforts illuminate the diversity and overlapping features between ADHD and ASD and offer a generalizable methodology for delineating transdiagnostic neurobiological variation in psychiatric populations.

## Methods

### Participants and Data Acquisition

#### Datasets

The ADHD-200, ABIDE-I, and ABIDE-II cohorts were utilized for our study. The ADHD-200 cohort^26^ includes 830 participants recruited from seven different scan sites. To facilitate independent validation of experimental findings, we split the whole dataset into a discovery and a replication sets. The ADHD-discovery set was formed using data from the three sites, including Kennedy Krieger Institute (KKI), Peking University (PKU), and New York University Child Study Center (NYU), since they provided a relatively balanced patient-control ratio in sample size (268 ADHD patients and 310 neurotypical controls). The ADHD-validation set was formed by 45 ADHD patients and 207 neurotypical controls from the remaining four sites: NeuroIMAGE Sample, Oregon Health and Science University (OSHU), University of Pittsburgh, and Washington University in St. Louis. The ABIDE cohort^27,28^ contains two independent datasets (ABIDE-I and ABIDE-II). The ABIDE-I dataset including 531 ASD patients and 571 neurotypical controls from 20 different sites was combined with the ADHD-discovery set for our discovery analysis of transdiagnostic subtypes. The ABIDE-II dataset including 453 ASD patients and 537 neurotypical controls from 16 different sites was combined with the ADHD-validation set to validate the generalizability of our subtype findings.

#### Phenotypical assessments

In the ADHD-200 cohort, phenotypical measures including FIQ, PIQ, and VIQ were assessed on one of the following scales: Wechsler Intelligence Scale for Children, Fourth Edition (WISC-IV), Wechsler Abbreviated Scale of Intelligence (WASI), and Wechsler Intelligence Scale for Chinese Children-Revised (WISCC-R). In the ABIDE cohort, FIQ scores are assessed based on one of the following scales: Differential Ability Scales II - School age (DAS-II), WASI, WISC, Wechsler Adult Intelligence Scales (WAIS), Hamburg-Wechsler Intelligence Test for Children (HAWIK-IV), and Wortschatztest(WST). VIQ scores are measured using DAS-II, WASI, WISC, WAIS, Peabody Picture Vocabulary Test (PPVT) and Stanford Binet (STANFORD). PIQ scores are evaluated by referring to DAS-II, WASI, WISC, WAIS, STANFORD and Raven’s Standard Progress Matrices (RAVENS).

#### Functional MRI acquisition and preprocessing

For both the ADHD-200 and ABIDE cohorts, rsfMRI data were collected with varying protocols and scanner parameters specific to each of the study sites. Details of imaging parameters can be found in literature^26–28^. All available rsfMRI data were preprocessed using the well-established fMRIPrep pipeline^65^. The T1-weighted image was corrected for intensity non-uniformity and then stripped skull. Spatial normalization was done through nonlinear registration, with the T1w reference^66^. Using FSL, brain features such as cerebrospinal fluid, white matter, and grey matter were segmented from the reference, brain-extracted T1 weighted image^67^. The fieldmap information was used to correct distortion in low-frequency and high-frequency components of fieldmap. Then, a corrected echo-planar imaging reference was obtained from a more accurate co-registration with the anatomical reference. The blood-oxygenation-level-dependent (BOLD) reference was then transformed to the T1- weighted image with a boundary-based registration method, configured with nine degrees of freedom to account for distortion remaining in the BOLD reference^68^. Head-motion parameters (rotation and translation parameters of volume-to-reference transform matrices) were estimated with MCFLIRT (FSL). BOLD signals were slice-time corrected and resampled onto the participant’s original space with head-motion correction, susceptibility distortion’s correction, and then resampled into standard space, generating a preprocessed BOLD run in MNI152NLin2009cAsym space. Automatic removal of motion artifacts using independent component analysis (ICA-AROMA)^69^ was performed on the preprocessed BOLD time-series on MNI space after removal of non-steady-state volumes and spatial smoothing with an isotropic Gaussian kernel of 6 mm FWHM (full-width half-maximum).

### Functional connectivity calculation and brain graph construction

With the preprocessed BOLD signals, we extracted regional time series based on the Schaefer parcellation^70^ of 100 regions of interest (ROIs). For each subject, functional connectivity was then calculated between these regional time series using Pearson’s correlation and partial correlation, respectively. We regressed out mean framewise displacement from functional connectivity features to alleviate head motion effects and then applied the well-established statistical harmonization tool ComBat ^29^ to reduce site effects. We included six personal characteristic data (PCD) comprising gender, age, handedness, and three IQ measures (FIQ, VIQ, PIQ) along with functional connectivity for the subsequent graph construction. Following literature^21,71^, each subject’s brain graph input is defined as *G* = (*V, E*) where node features for each ROI is *V_i_∈* R*^ROIs^*^+^*^PCD^* = R^106^which concatenates the *i*-th row of the Pearson’s correlation matrix with the corresponding PCD. The edge feature *E_i j_ ∈* R indicates the partial correlation connectivity between ROI *i* and *j*. A range of sparsity values are tested on the partial correlation and the optimal sparsity is set as 0.6 to trade off between efficiency and precision. As contrastive learning requires multiview inputs from the same subject to form a positive pair, we generate three graphs for each subject using either multiple scans of the same subject or three non-overlapping ROI slices if the subject only had a single scan.

### Contrastive graph representation learning

Our proposed contrastive representation framework consists of two components (Supplementary Figure 1): contrastive graph learning (CGL) and dynamic graph classification (DGC). CGL aims to train a self-supervised neural network that can represent high dimensional subject features. DGC utilizes diagnostic label information to further improve the representation by quantifying patient-specific variations.

The motivation behind CGL is that two functional connectivity views coming from the same subject should form a “homogeneous” pair whereas different patient views should form a “heterogeneous” pair. More specifically, we can treat multiple fsfMRI slices obtained from one individual as different views of such subject, and therefore the functional connectivity extracted by those slices should share higher similarities than two random functional connectivity matrices chosen from two different subjects. Such inductive bias avoids the adoption of labels to train deep networks, which effectively alleviates overfitting and has been testified in many scenarios^72^. For implementation, the brain graph *G* was firstly passed to several spectral graph convolutional networks^73^ defined as:

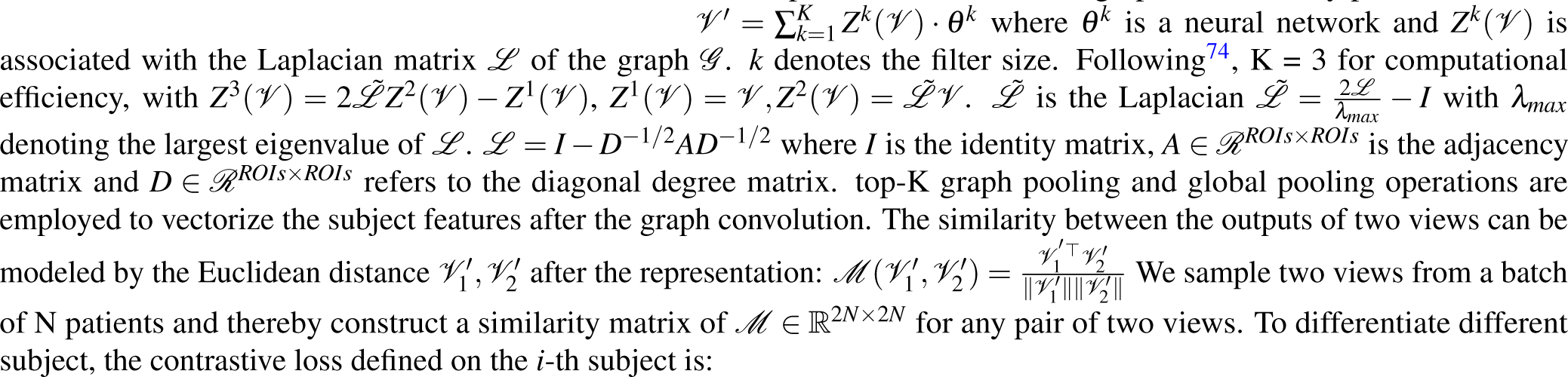

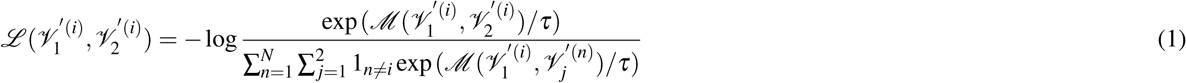

where *τ* is a temperature factor to control the desired attractiveness strength and 1(*·*) = *{*0, 1*}*. The numerator aims to enhance homogeneous attraction whereas the denominator minimize the overall heterogeneous similarities across the population. The overall loss function is aggregated over all the pairs: 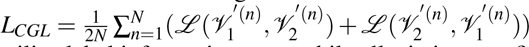 The resulting representations of patients *V ^′^* are then fed to the DGC to utilize label information meanwhile alleviating overfitting.

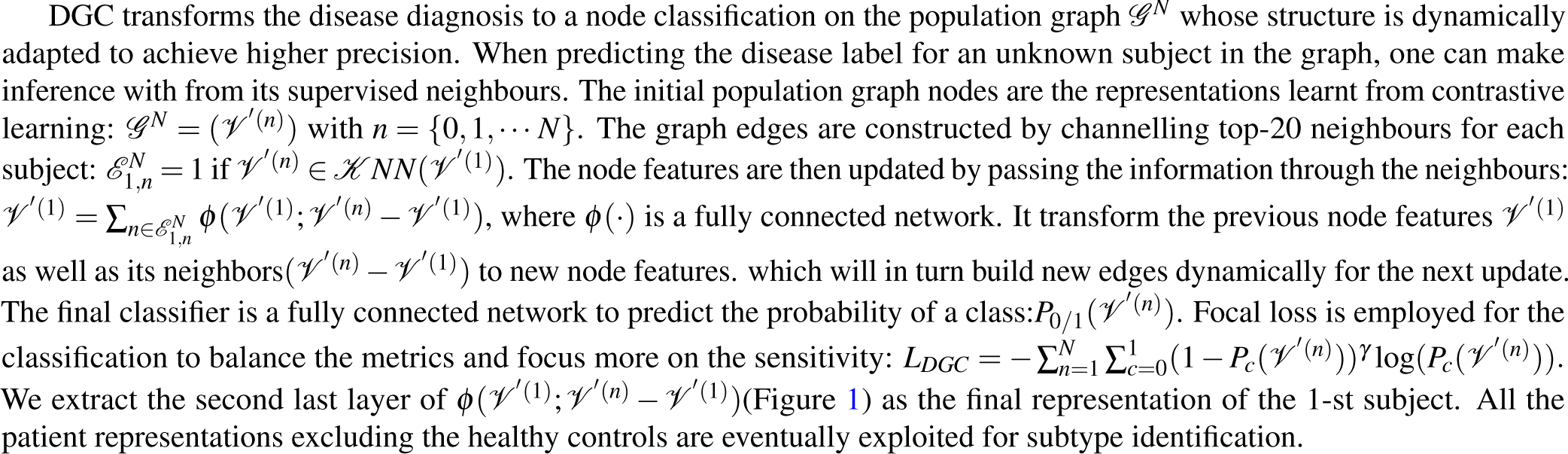

### Subtype identification

After acquiring patient representation with the label-aimed contrastive learning, we apply spectral clustering on the population graph 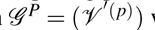 with *p* = *{*0, 1, · · · P} denoting all the patients. Spectral clustering can explicitly model patient similarities and offer the customized cluster count by selecting the top-*K* eigenvectors. Given the Laplacian matrix of 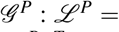 *I − D^−^*^1^*^/^*^2^*AD^−^*^1^*^/^*^2^, spectral clustering firstly computes its eigenvalues and eigenvectors: 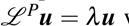 where 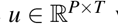 with *T ≤ P* indicates the eigenvectors and *λ* is the eighvalue. We rank the eigenvalues and choose top-*K* corresponding eigenvector 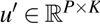 as a low-dimensional orthogonal mapping of the patients where a K-means is applied. The clustering labels 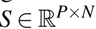 are mapped back to the patients as subtype results. The optimal number of subtypes was selected based on the highest Calinski-Harabasz index.

### Discriminative functional connectivity identification through partial derivatives

The subtypes are observed on the representation space 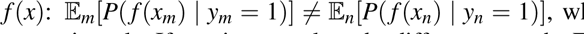, where two communities M and N from the disease group do not perform consistently. If we aim to analyse the differences on the FC inputs, which can showcase medical significance and interpretation, the gradient-based method offers a promising solution. It has been shown to demonstrate feature importance, also known as the saliency map^75,76^. The motivation is that suppose a prediction *y* = *f* (*x*) can be approximated with a linear function *y* = *Wx* + *b*, then the partial derivative *∂y/∂x* = *W* showcases the crucial inputs with the greatest influence on the outputs. Based on this, we train a subtype classifier using the subtyping of the training set as the ground-truth. The classifier replaces the last layer of the original contrastive framework with a new subtyping classifier. The reason for such layer modification rather than utilizing the gradients of original framework is the shift of focus from the disease to the subtypes. The gradients of the original disease classifier can only indicate the connectivities contributing to the disease, which would present spurious similarities between both subtypes and fail to showcase the subtyping differences. Finally, the gradients of the subtyping outputs over the connectivity inputs including both the partial and ’s connectivities are derived to highlight important connectivities.

## Data availability

The data supporting the results in this study are available within the paper and its Supplementary Information. The neuroimaging datasets are publicly available from ADHD-200 (http://fcon_1000.projects.nitrc.org/indi/adhd200/), ABIDE-I and ABIDE-II (https://fcon_1000.projects.nitrc.org/indi/abide/).

## Code availability

The custom code used in this study is available for research purpose from the corresponding author upon reasonable request.

## Acknowledgements

Y.Z. was supported by NIH grant nos. R21MH130956, R01MH129694, and R21AG080425, and Lehigh University FIG (FIGAWD35), CORE, and Accelerator grants. G.A.F. was supported by NIH grant nos. R01MH132784 and R01MH125886 and grants from the Brain and Behavior Research Foundation and One Mind – Baszucki Brain Research Fund. T.D.S. was supported by NIH grant nos. R37MH125829, R01EB022573, R01MH112847, and R01MH120482.

## Supplementary

**Figure S1.**
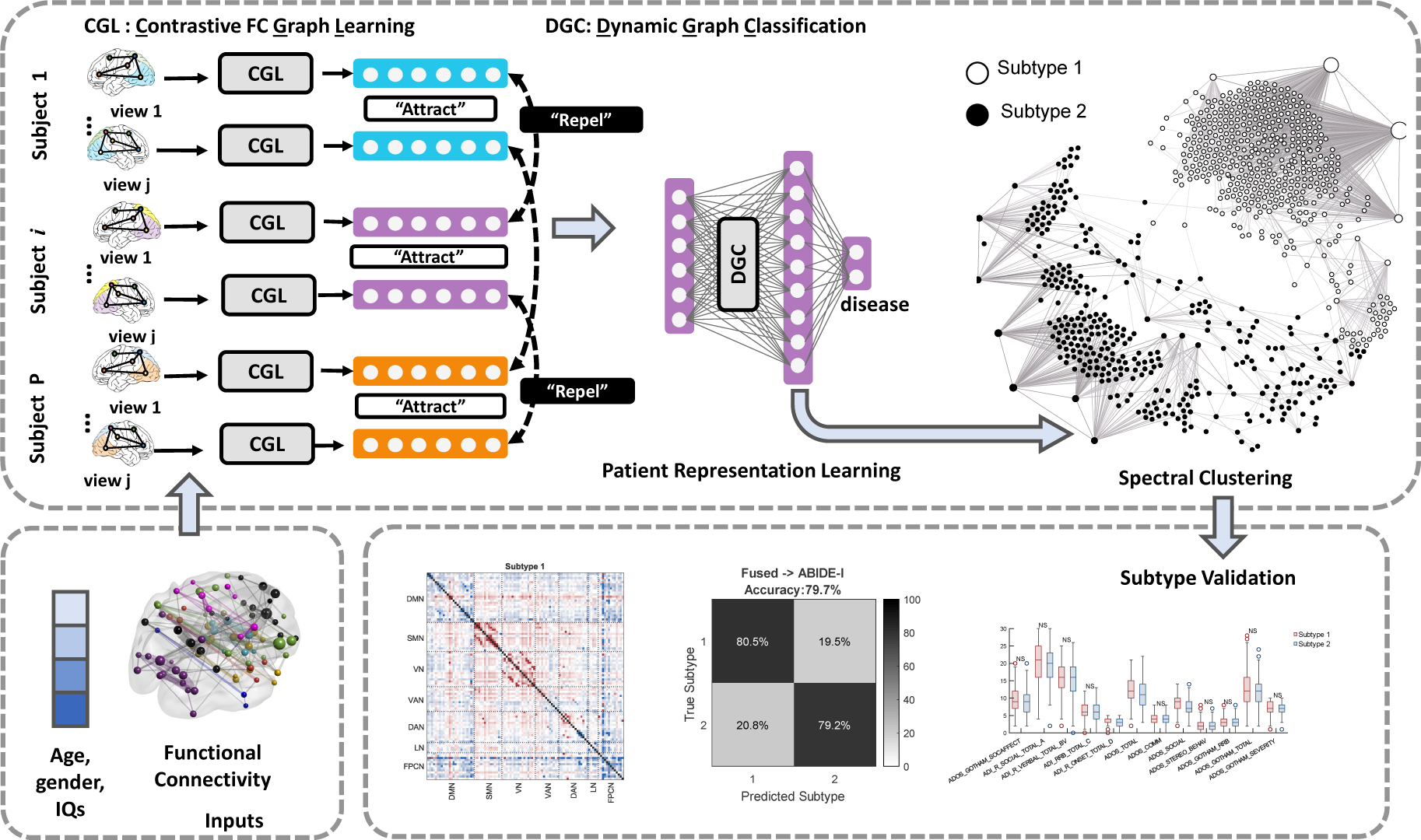
Illustration of contrastive learning framework for transdiagnostic subtype identification.

**Figure S2.**
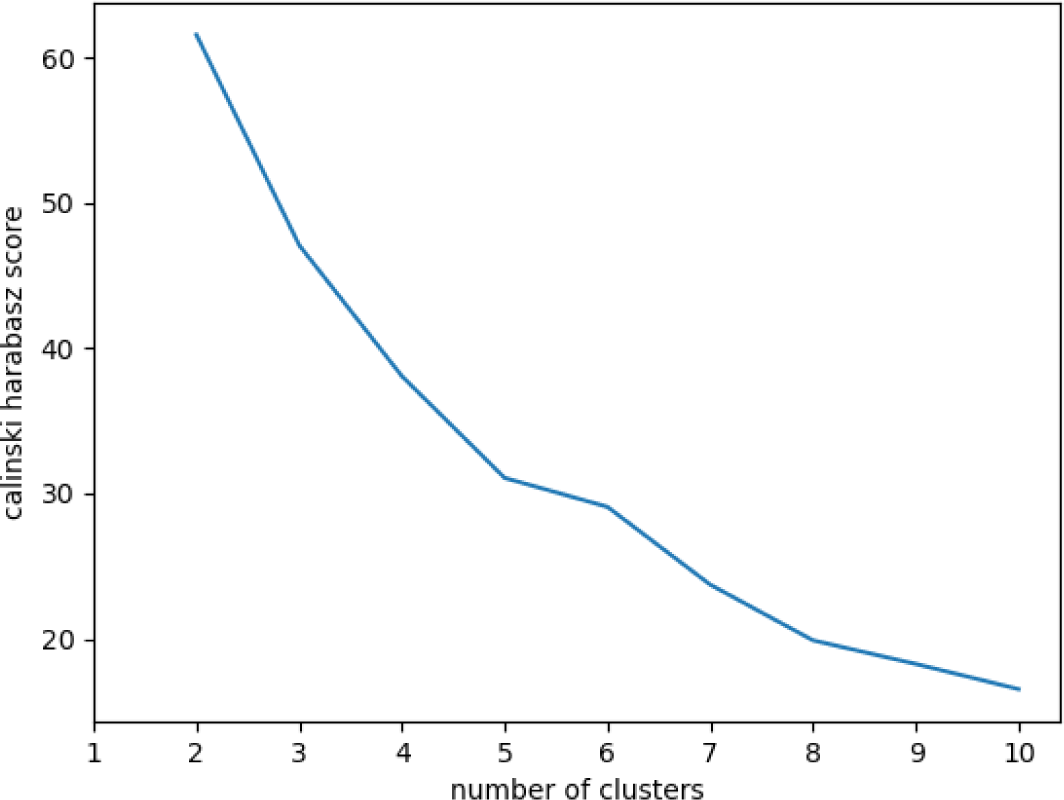
Selection of the optimal number of clusters. We run the spectral clustering method on the transdiagnostic ASD and ADHD population. We set varying number of clusters ranging from 2 to 10 for the clustering algorithm whose input is a distance matrix between subjects. For each choice of the number we compute the corresponding Calinski-Harabasz score that measures the ratio of inter-cluster distance and intra-cluster distance. Therefore the highest Calinski-Harabasz score suggests clear-defined clusters.

**Figure S3.**
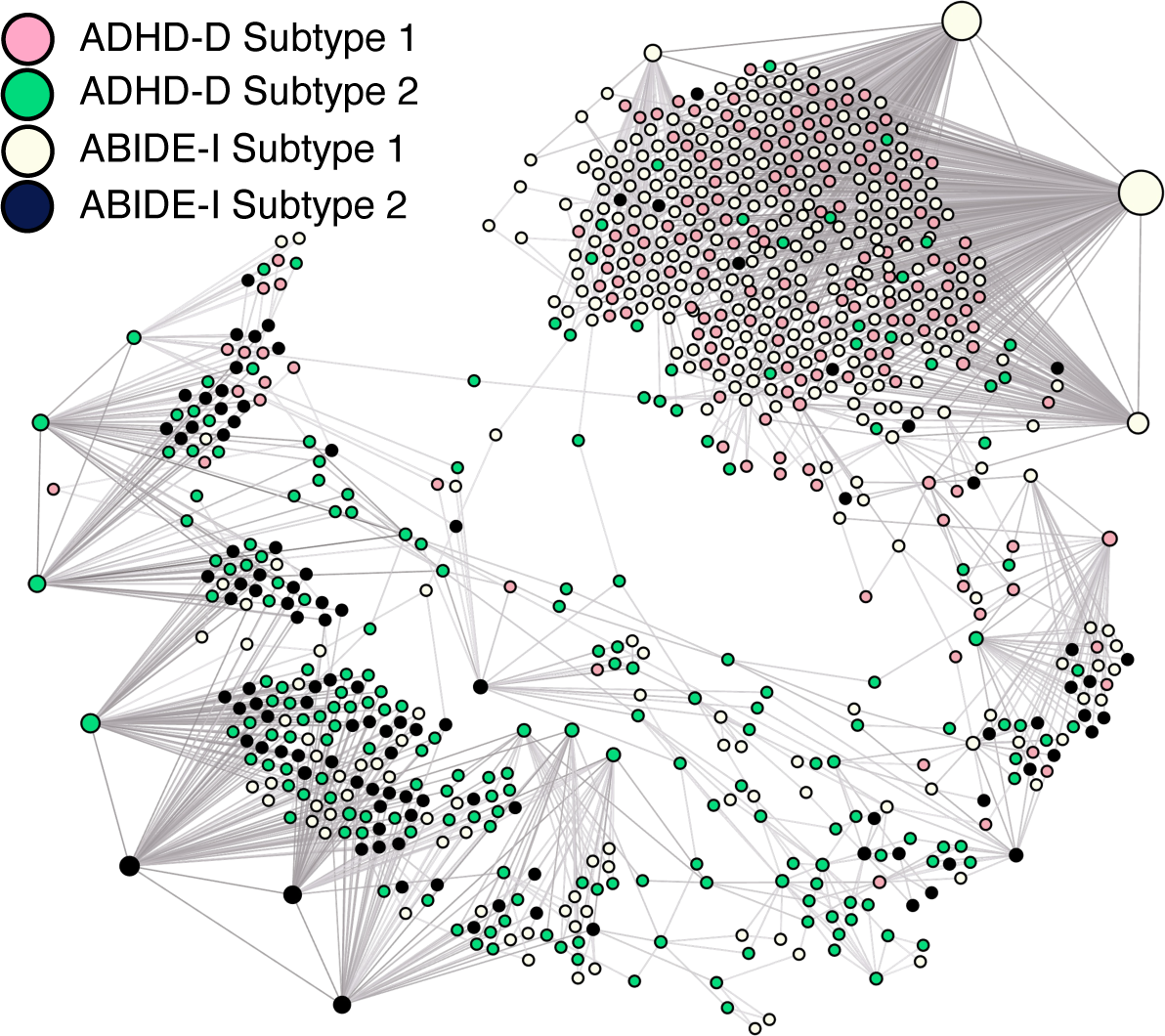
Subtype relationships between the individual disease. Two subtypes are obtained separately in ABIDE-I and ADHD-discovery set (ADHD-D) using the same pipeline to showcase transdiagnostic heterogeneity and comorbidity. The population graph was generated by the transdiagnostic discovery set but the node labels are obtained through clustering on individual diseases. Their results are mapped to the transdiagnostic population graph with different colors indicating the subtype labels acquired from individual diseases.

**Figure S4.**
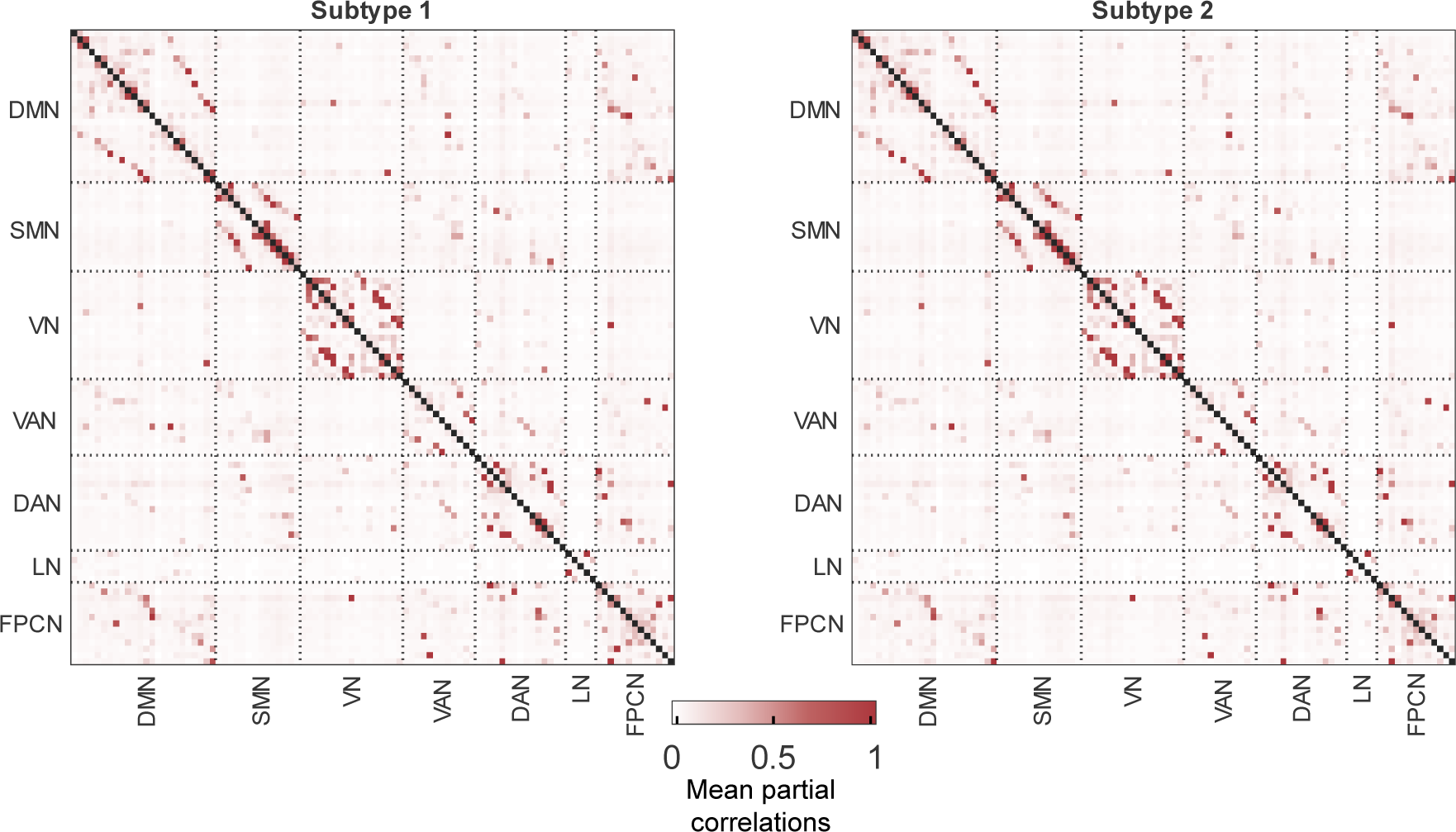
**Mean connectivity inputs of the two subtypes extracted from the transdiagnostic set of ABIDE-I and ADHD discovery set**. The color bar indicates the mean partial connectivity values.

**Figure S5.**
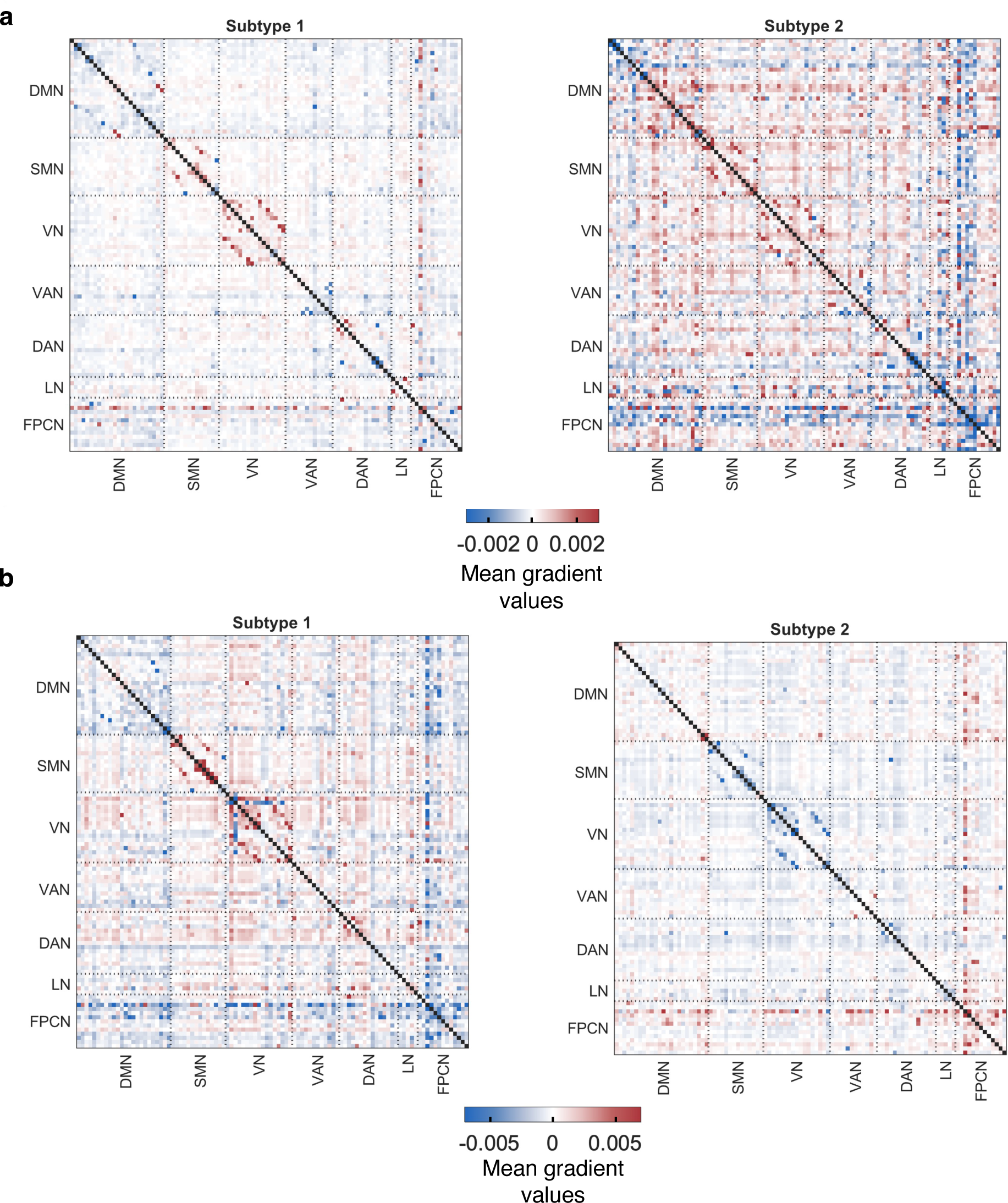
Connectivity gradients of two subtypes obtained from the individual discovery sets. The gradients are obtained from running the subtyping on individual sets separately, where positive gradients mean the positive contribution of a connectivity to the subtype. **a**. Mean connectivity gradients of the two subtypes on ABIDE-I. **b.** Mean connectivity gradients of the two subtypes on the ADHD-discovery set.

**Figure S6.**
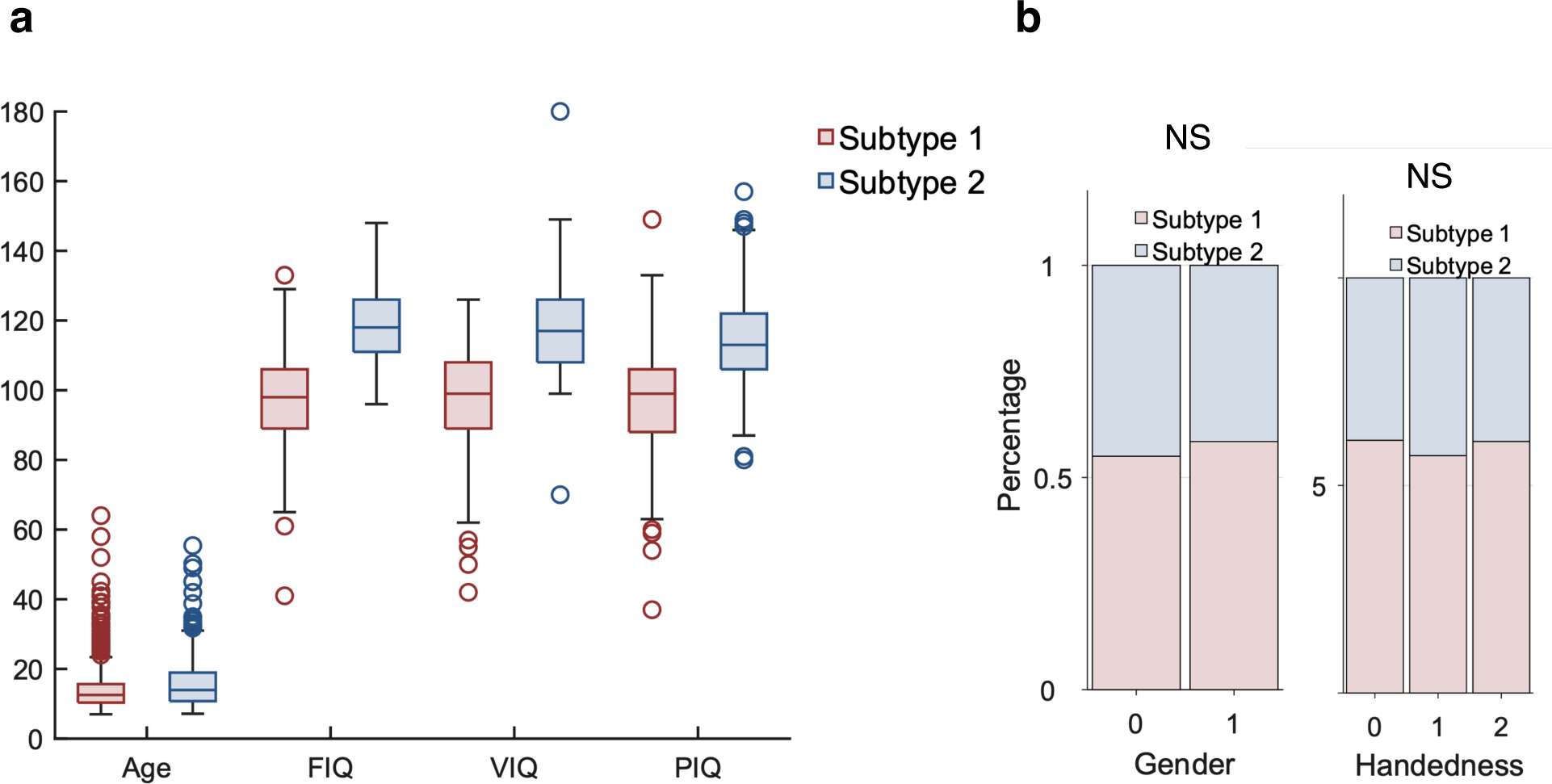
Demographic and clinical differences on the transdiagnostic discovery set (ABIDE-I + ADHD discovery set). **a.** Boxplot for numerical features including age and three IQ cognitive measurements,where ‘NS’ indicates non- significant t-test results. **b.** Stacked percentage barplot for categorical features that showcases the percentage of the two subtypes in each feature category, such as gender (0: Female, 1: Male) and handedness (0: Left, 1: Right, 2: Ambidextrous. Continuous values between 0 to 1 in ADHD are categorized as Ambidextrous). ‘NS’ shows non-significant chi-square test results.

**Figure S7.**
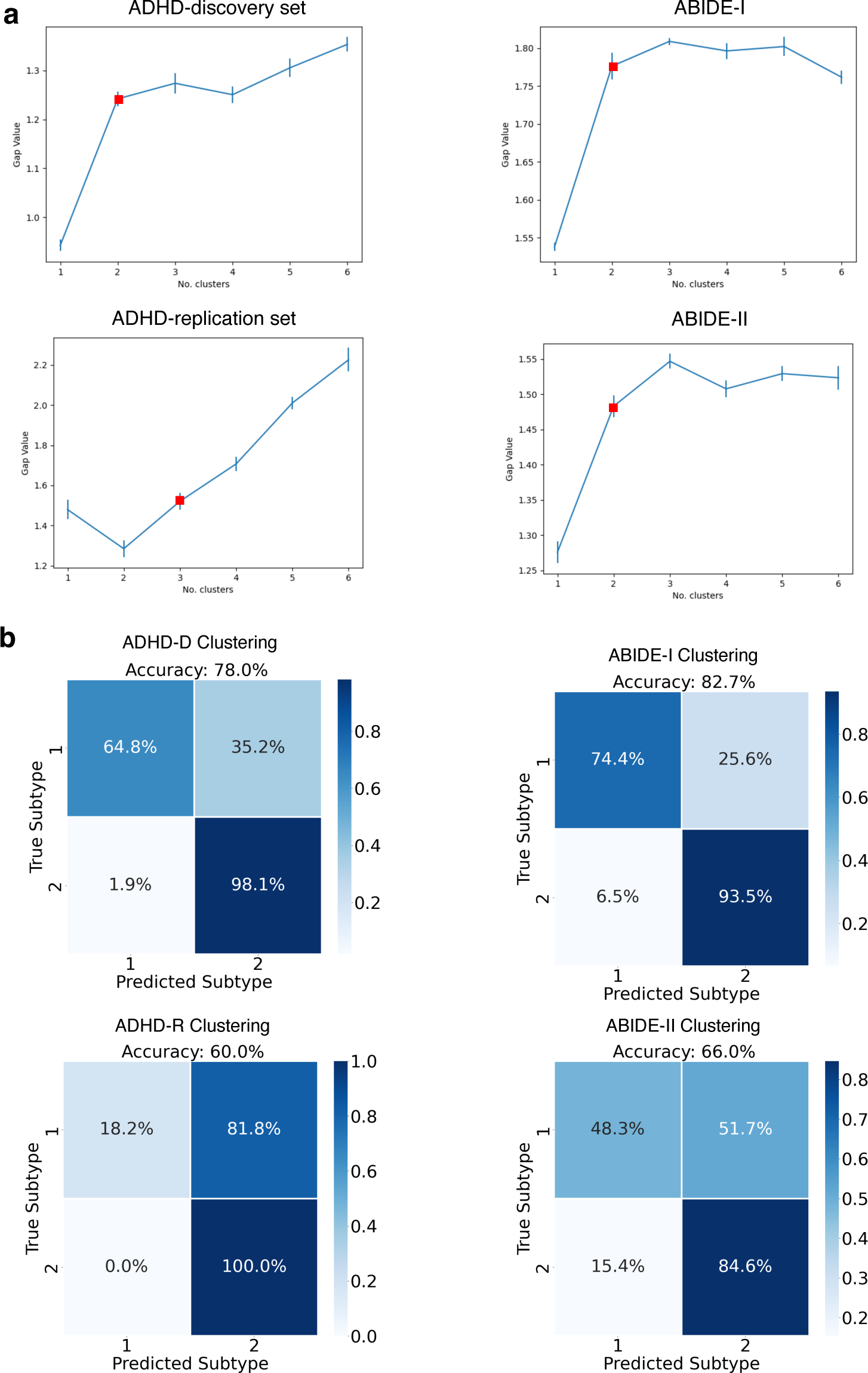
Clustering based on demographic information and cognitive measures for the discovery and replication datasets. Six features including Gender, Handedness, Age, FIQ, VIQ, PIQ are utilized to train the subtyping cluster. **a.** The optimal number of clustering (noted with red squares) found in 2 discovery and 2 replication sets using gap statistic criterion. The error bars indicate standard deviation. **b.** Confusion matrices of comparing the subtyping labels obtained by our pipeline (ground truths) and subtyping (predictions) using biological features as such age, gender are shown with confusion matrices.

**Figure S8.**
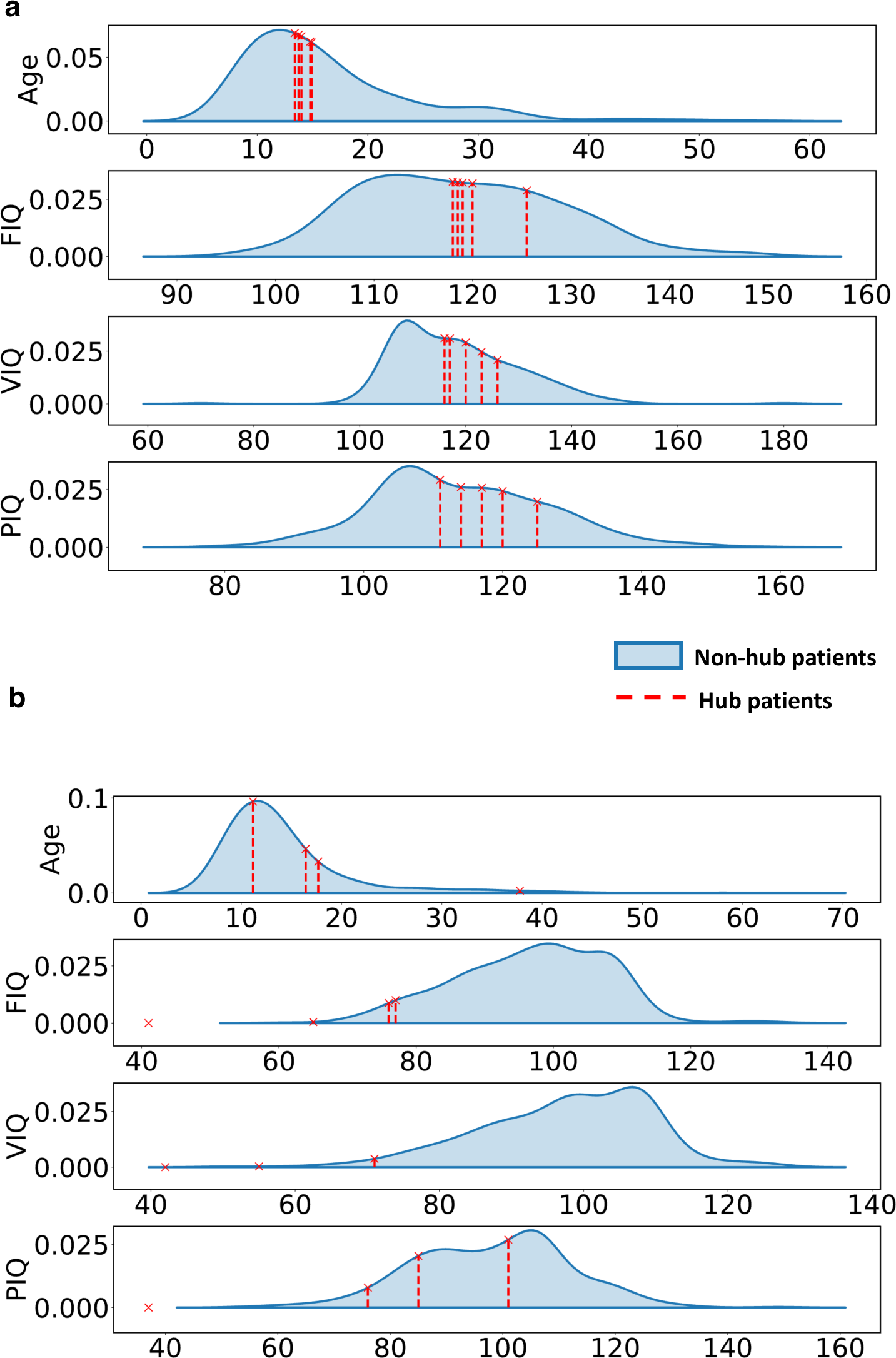
PCD differences between the hub and non-hub patients in the transdiagnostic discovery set. Hub patients are defined by the patients with the highest node degree in the population graph. We estimate the distributions of the age and three cognitive IQs for the non-hub patients in each subtype. Their estimates on the hub data are shown in crosses and the dashed line, where a long dashed line indicate normality and small value indicates abnormality. **a.** Six hub patients verus the rest non-hub patients in subtype 1. The non-hub distribution is fitted by the Gaussian kernel density estimation. **b.** Four hub patients versus the rest non-hub patients in subtype 2.

**Figure S9.**
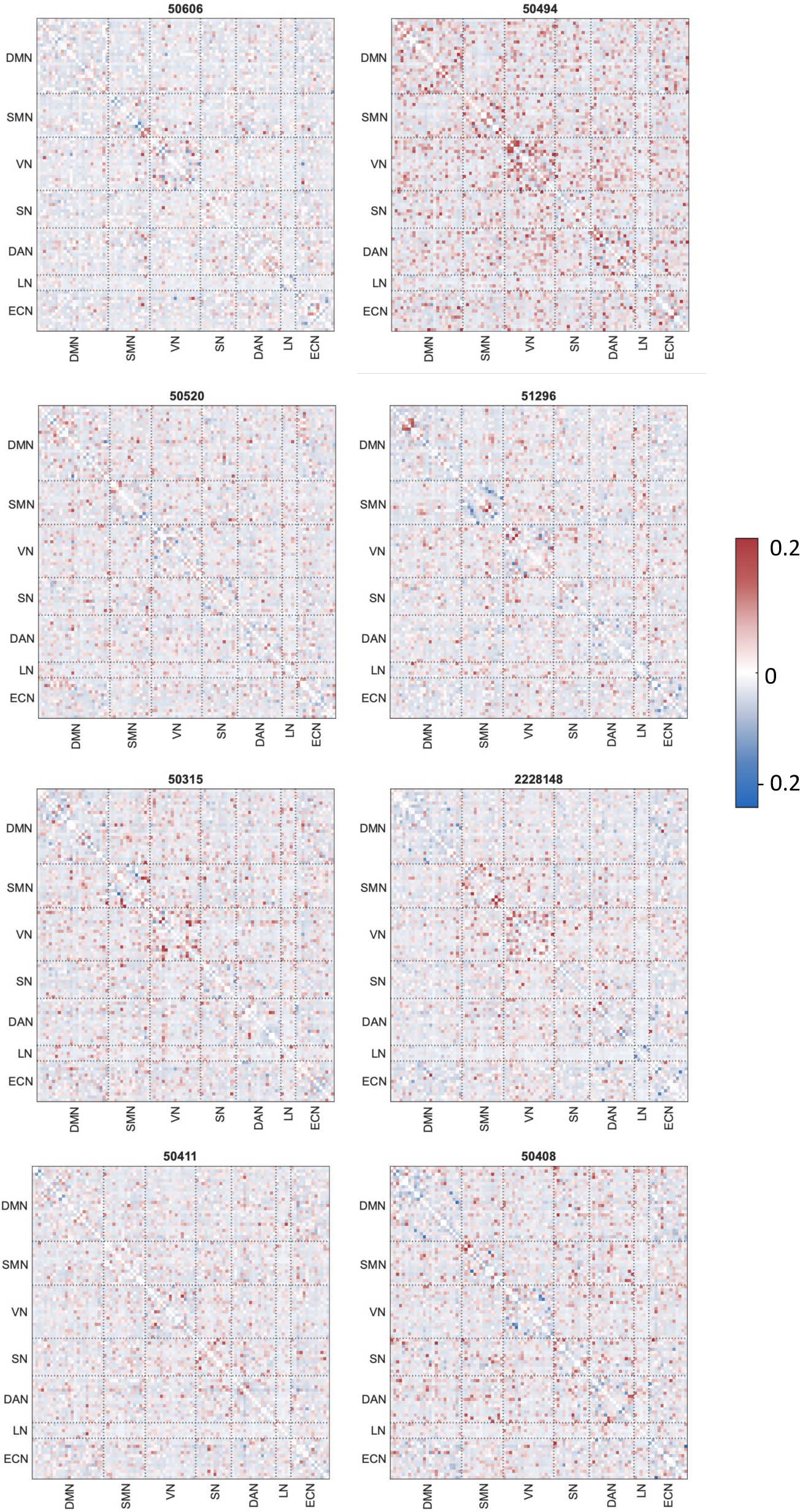
FC differences between the hub and non-hub patients in the transdiagnostic discovery set. The label indicate the names of the hub subjects. The matrix values show the different between the FC inputs of a hub patient and the mean FC values from the non-hub patients within its corresponding subtype category.

**Table S1.**
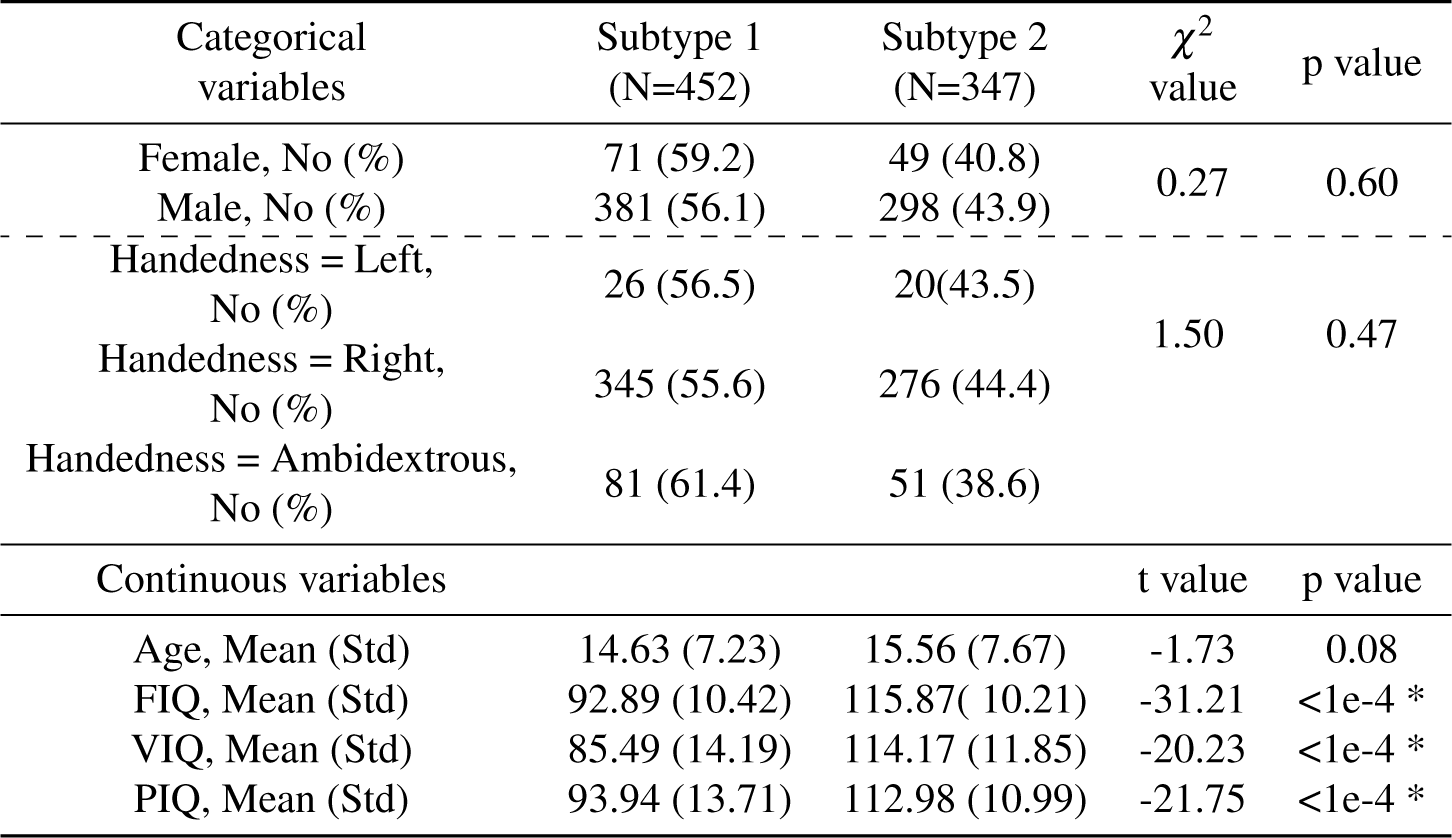
Demographic and clinical characteristic differences between the two subtypes on the transdiagnostic dataset of ADHD-discovery set + ABIDE-I. * indicates significant differences with p value < 0.05

**Table S2.**
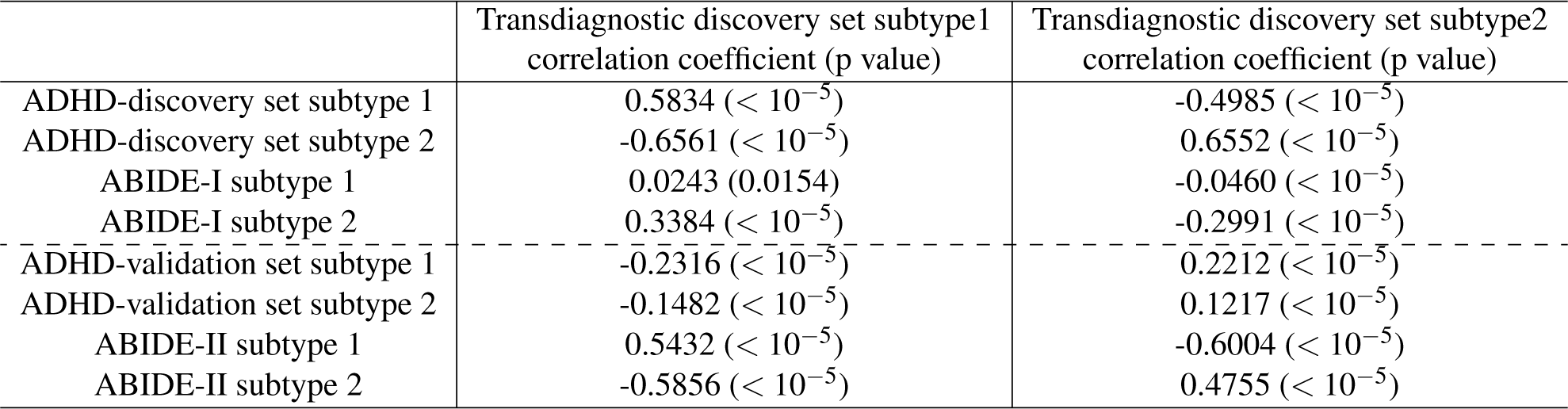
Similarity measure between individual dataset and the transdiagnostic discovery set using correlation coefficient. * indicates significant differences with p value < 0.05

**Table S3.**
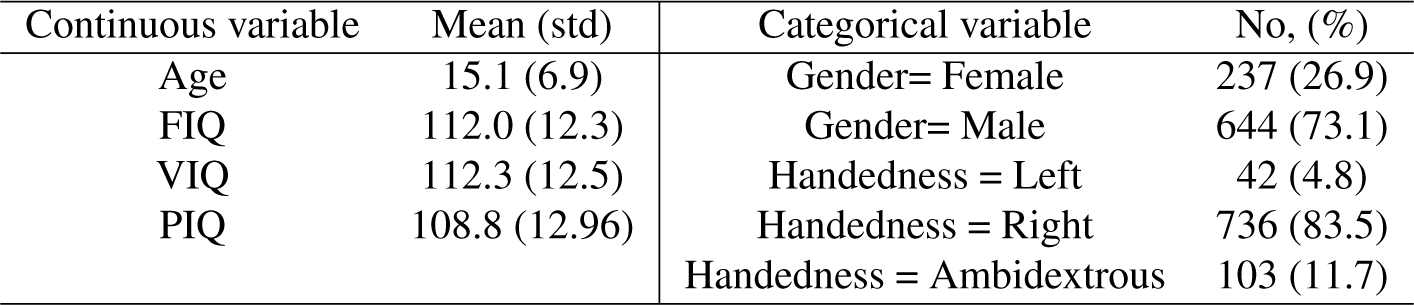
Demographic characteristics and cognitive assessments of healthy controls from the transdiagnostic discovery set. * indicates significant differences with p value < 0.05

**Table S4.**
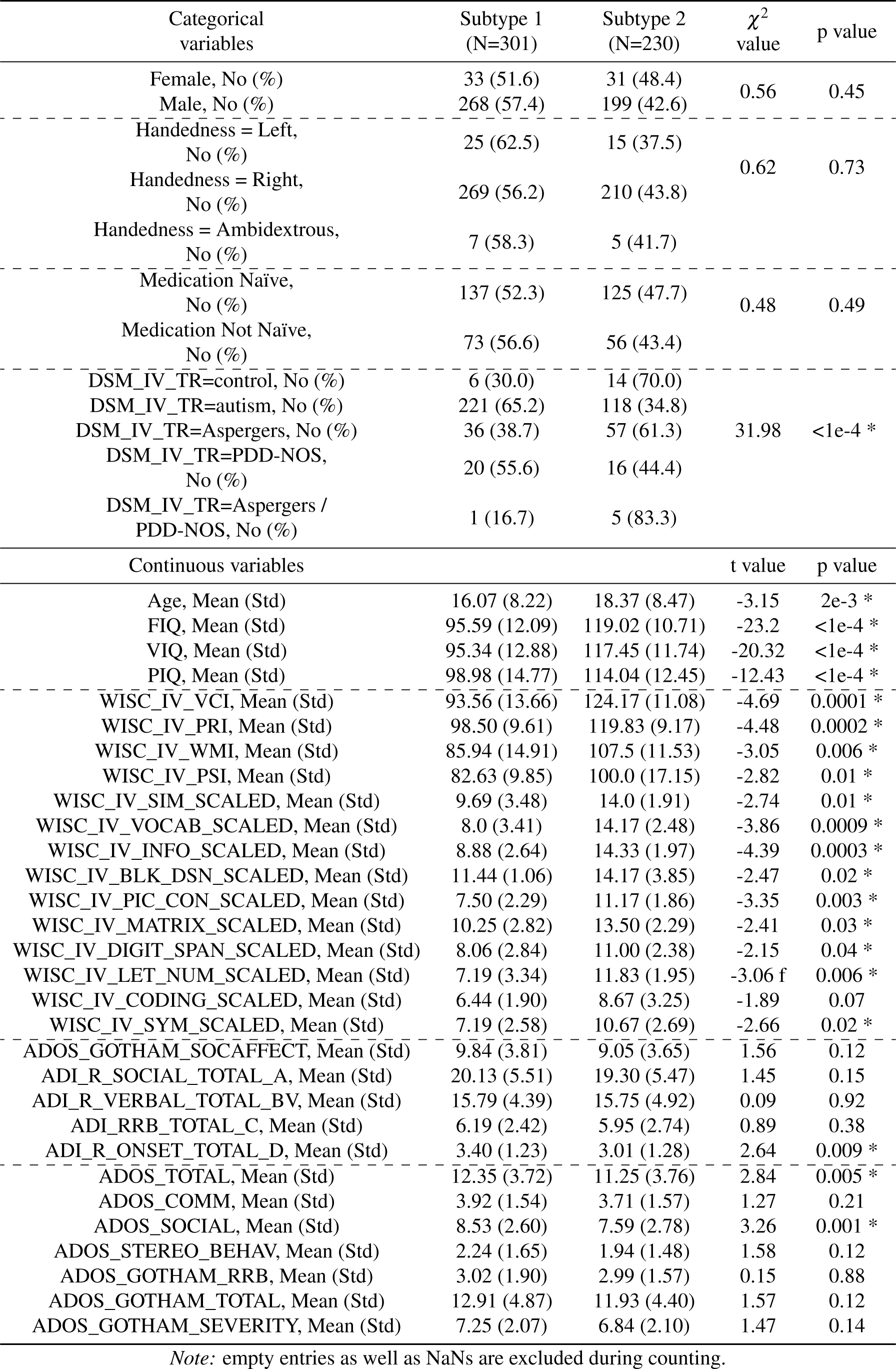
Demographic characteristics and clinical variables between the two subtypes on the ABIDE-I dataset.

**Table S5.**
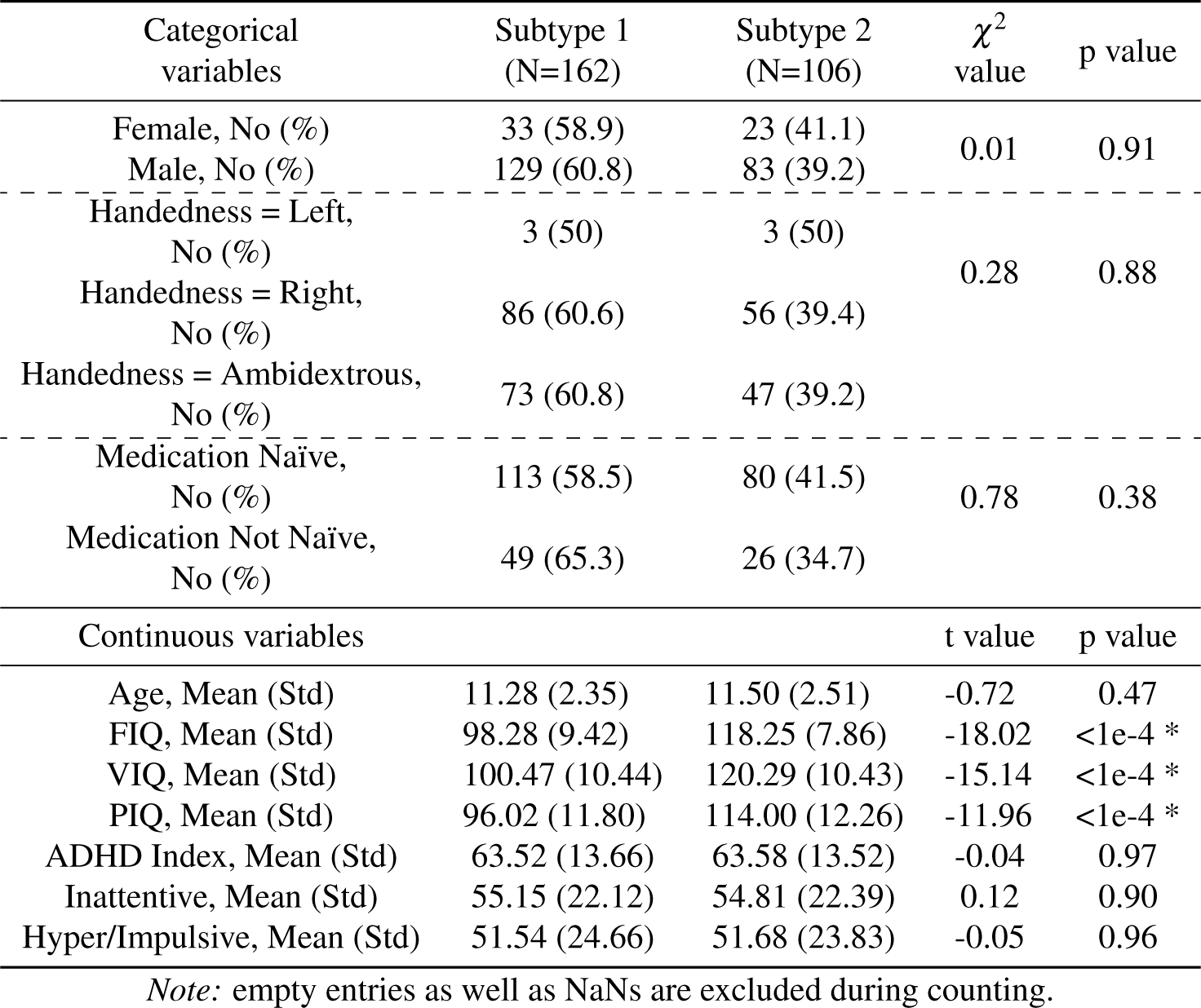
Demographic characteristics and clinical variables between the two subtypes on the ADHD-discovery set. The subtyping labels are obtained from the transdiagnostic discovery set. * indicates significant differences with p value < 0.05

**Table S6.**
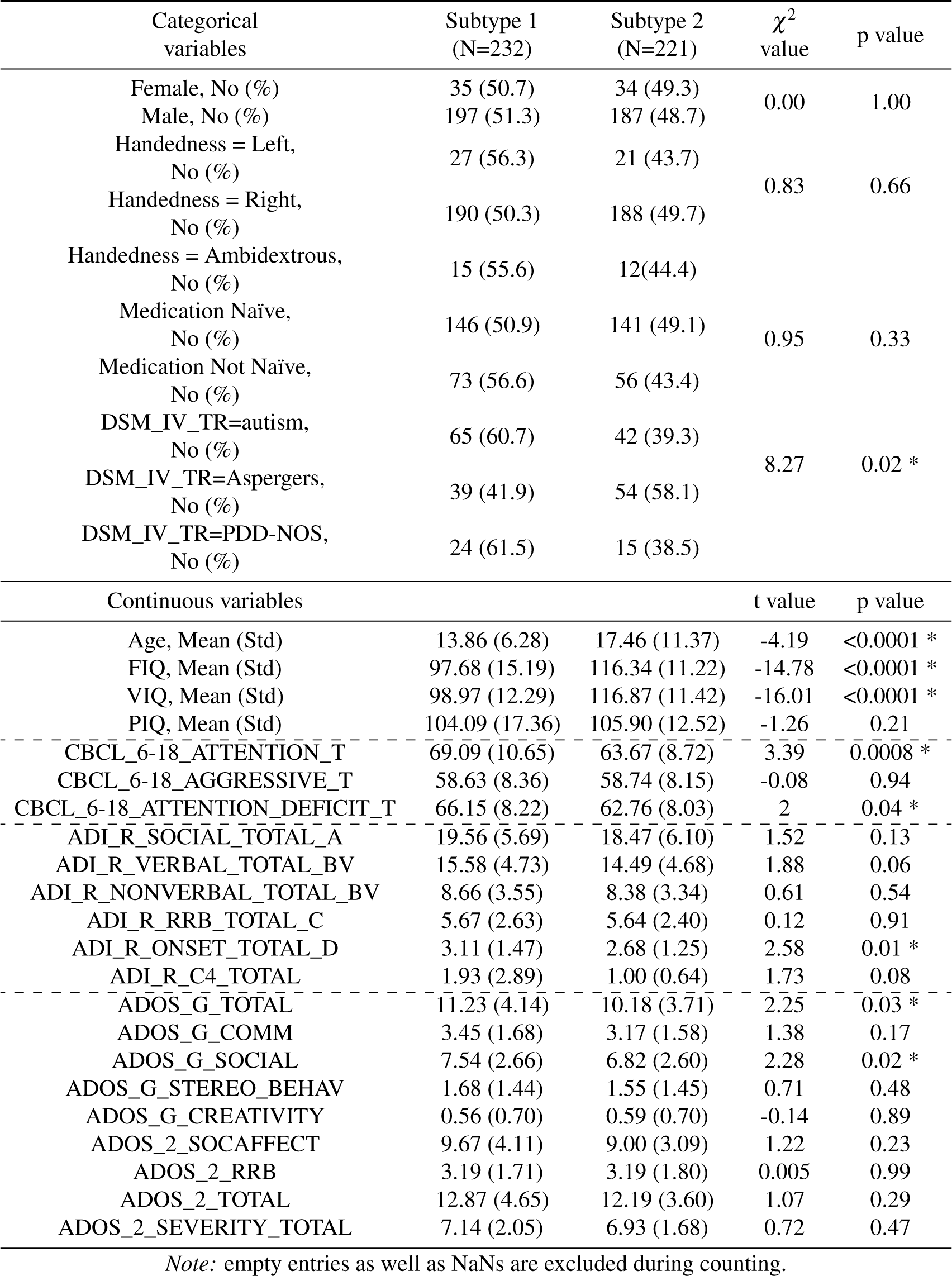
Demographic characteristics and clinical variables between the two subtypes on the ABIDE-II dataset. * indicates significant differences with p value < 0.05

**Table S7.**
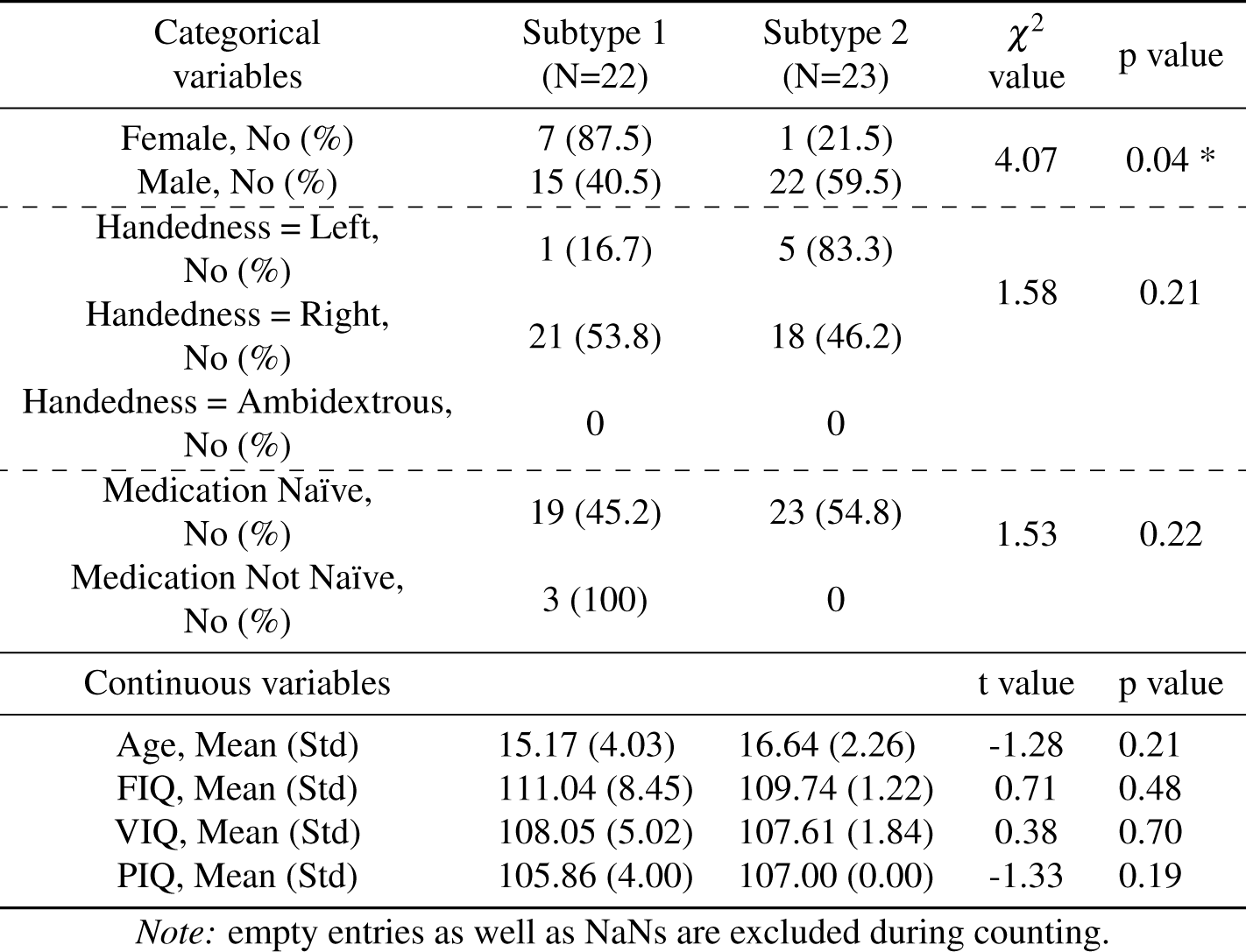
Demographic characteristics and clinical variables between the two subtypes on the ADHD-validation set. * indicates significant differences with p value < 0.05

## Notes

### Competing Interest Statement

The authors have declared no competing interest.

